# POSTRE: a tool to predict the pathological effects of human structural variants

**DOI:** 10.1101/2022.06.20.496902

**Authors:** Víctor Sánchez-Gaya, Álvaro Rada-Iglesias

## Abstract

Understanding the pathological impact of non-coding genetic variation is a major challenge in medical genetics. Accumulating evidences indicate that a significant fraction of genetic alterations, including structural variants (SVs), can cause human disease by altering the function of non-coding regulatory elements, such as enhancers. In the case of SVs, described pathomechanisms include changes in enhancer dosage and long-range enhancer-gene communication. However, there is still a clear gap between the need to predict and interpret the medical impact of non-coding variants, and the existence of tools to properly perform these tasks. To reduce this gap, we have developed POSTRE (Prediction Of STRuctural variant Effects), a computational tool to predict the pathogenicity of SVs implicated in a broad range of human congenital disorders. By considering disease-relevant cellular contexts, POSTRE identifies SVs with either coding or long-range pathological consequences with high specificity and sensitivity. Furthermore, POSTRE not only identifies pathogenic SVs, but also predicts the disease-causative genes and the underlying pathological mechanism (e.g, gene deletion, enhancer disconnection, enhancer adoption, etc.). POSTRE is available at https://github.com/vicsanga/Postre.

## Introduction

Structural Variants (SVs) represent one of the greatest sources of genetic variation in the human genome (Feuk et al., 2006; Stankiewicz and Lupski, 2010). This group of genetic alterations includes: deletions, duplications, inversions, insertions, and translocations. Their size ranges from a few base pairs (∼50) to several megabases (Mbs) (Ho et al., 2020). SVs can cause phenotypic diversity and a broad set of human disorders, including congenital abnormalities and various cancer types (Spielmann et al., 2018). The pathogenicity of SVs can be triggered by two main mechanisms: (i) direct effects on genes (e.g. “coding” pathomechanisms: changes in gene dosage, gene truncations, formation of fusion transcripts) and (ii) changes in the non-coding regulatory landscape of disease relevant genes that alter their expression levels (i.e. “long-range” pathomechanisms) (Spielmann et al., 2018; Sánchez-Gaya et al., 2020; Krude et al., 2021). However, it is currently challenging to predict or interpret the functional consequences of SVs. This is particularly true for SVs that affect gene expression through long-range regulatory mechanisms (Spielmann et al., 2018; Lappalainen et al., 2019; Krude et al., 2021) due to the limited understanding of the non-coding regulatory genome and the cell-type specific functions of distal regulatory elements, such as enhancers.

The vast majority (>90%) of disease associated variants are located within the non-coding fraction of the human genome (Maurano et al., 2012; Krijger and De Laat, 2016, Zhu et al., 2017; Elgar and Vavouri, 2008), mainly within putative enhancers (Krijger and De Laat, 2016). Enhancers are important and abundant cis-regulatory elements (Ong and Corces, 2011; Wittkopp and Kalay, 2012) that positively control the expression of their target genes in space and time, and are major determinants of cell-type specific gene expression programs (Wray, 2007; Bulger and Groudine, 2010, 2011; Buecker and Wysocka, 2012). Enhancers have been globally identified in hundreds of human cell types and tissues using universal epigenomic approaches (Heintzman et al., 2009; Creyghton et al., 2010; Rada-Iglesias et al., 2011). Enhancers can control gene expression over large genomic distances (>1Mb) (Lettice, 2003; Sagai et al., 2005; Long et al., 2020), often skipping proximal genes while controlling the expression of more distally located ones (Sanyal et al., 2012). Consequently, linking enhancers with their target genes is not an obvious task. In this regard, 3D genome organization studies based on Hi-C technology (Lieberman-Aiden et al., 2009) revealed that genomes are organized in large (Mb-scale) self-interacting domains often referred to as topologically associating domains (TADs) (Dixon et al., 2012). TADs represent fundamental regulatory domains as (i) they facilitate enhancer–gene interactions within them (Dixon et al., 2012; Nora et al., 2013; Rao et al., 2014; Spielmann et al., 2018) and (ii) insulate genes from contacting ectopic enhancers located in other TADs (Lupiáñez et al., 2016). SVs can disrupt TAD organization, which in turn can rewire enhancer-gene communication and lead to pathological changes in gene expression (Lupiáñez et al., 2016; Spielmann et al., 2018; Sánchez-Gaya et al., 2020). This can occur through two alternative long-range regulatory mechanisms: (i) SVs can lead to gene silencing (loss of function (LOF)) by disconnecting genes from their cognate enhancer/s (i.e. enhancer disconnection) (Laugsch et al., 2019), or through the deletion of enhancers (i.e. enhancer deletion) (Benko et al., 2009); (ii) SVs can lead to gains in gene expression (gain of function (GOF)) by enabling enhancers to interact with non-target genes (i.e. enhancer adoption/hijacking) (Lupiáñez et al., 2015), or by the duplication of enhancers (Lohan et al., 2014). Nevertheless, predicting the pathological consequences of TAD disruption is complicated by the fact that, besides TAD organization, other genetic and epigenetic features also contribute to productive gene-enhancer communication (Ghavi-Helm, 2019; Sánchez-Gaya et al., 2020; Pachano et al., 2021; Batut et al., 2022; Bergman et al., 2022; Zuin et al., 2022).

Since gene expression programs change in space and time, it is fundamental to assess the pathological impact of SVs in the relevant cellular contexts. For instance, a deletion in Chr17 causes a pathogenic downregulation of SOX9 in the neural crest but not in other cell types, such as embryonic stem cells (ESCs) or chondrocytes (Long et al., 2020). Importantly, this deletion eliminates enhancers that are specifically active in neural crest cells (NCCs) and that, consequently, control *SOX9* expression in this cell type but not in others. Additionally, the same SVs might differentially affect the expression of the same gene depending on the cellular context. For example, a deletion could eliminate enhancers in one cell type in which the gene is active and lead to gene silencing (pathogenic LOF), whereas in another cell type in which the same gene is inactive, the deletion could eliminate a TAD boundary and lead to gene overexpression through enhancer adoption mechanisms (pathogenic GOF) (Spielmann et al., 2018). Therefore, although the recurrency of non-coding SVs within specific genomic loci could help identifying the genes involved in certain disorders, the exact pathomechanisms can only be predicted and experimentally validated if cell type specific regulatory landscapes (i.e. enhancers and TADs) are considered. During the last few years, and partly due to the efforts of large international consortia, different types of genomic data (e.g. epigenomic, transcriptomic and Hi-C data) have been generated in hundreds of different human cell types and tissues (Dunham et al., 2012; Satterlee et al., 2019). Furthermore, most of this data is available through public databases such as GEO (Barrett et al., 2013). In principle, these genomic datasets could be integrated in order to predict the pathological consequences of SVs in a cell type-specific manner (Sánchez-Gaya et al., 2020). Nonetheless, finding the appropriate datasets for the disease of interest, as well as their subsequent processing, analysis and integration can be very time-consuming and may require advanced bioinformatic skills. Consequently, the diagnosis and interpretation of how SVs might cause human disease remains complicated and the pathogenicity of SVs identified in hundreds of patients is currently unknown (Federici and Soddu, 2020). To overcome these limitations is essential to implement user-friendly computational tools that can be used by a broad scientific community (Sánchez-Gaya et al., 2020; Krude et al., 2021).

Taking the previous challenges into account, we developed POSTRE (<u>P</u>rediction <u>O</u>f <u>STR</u>uctural variant <u>E</u>ffects), a computational tool that, thanks to its user-friendly graphical interface, facilitates the cell type-specific analysis of SVs implicated in congenital disorders. Compared to other SV analysis tools, POSTRE is able to predict both coding and long-range pathomechanisms in a cell type/tissue-specific manner. The current version of POSTRE can handle SVs potentially implicated in limb, craniofacial, cardiac and neurodevelopmental congenital abnormalities, thus covering a broad set of congenital diseases (Trainor, 2010; Jeste, 2015; Kirby, 2017; Hansen et al., 2018; Wu et al., 2020; Barik et al., 2021). Here we extensively describe how POSTRE works and illustrate how it can predict the pathological consequences of SVs, particularly those implying long-range regulatory mechanisms, with high sensitivity and specificity. POSTRE is available at https://github.com/vicsanga/Postre.

## Results

### POSTRE overview

POSTRE is a software developed to predict the pathological impact of SVs implicated in a broad set of congenital abnormalities (i.e. limb, craniofacial/head&neck, cardiac or neurodevelopmental). In comparison with previous tools (see POSTRE benchmarking section for more details), POSTRE can analyze SVs with direct effects on protein coding genes as well as SVs acting through long-range regulatory mechanisms that alter gene expression. Furthermore, POSTRE can analyze not only Copy Number Variants (CNVs) (i.e. deletions and duplications) (Hertzberg et al., 2022), but also balanced inversions and translocations. In this section we provide an overview of the tool (Figure 1a-c).

**Figure 1.**
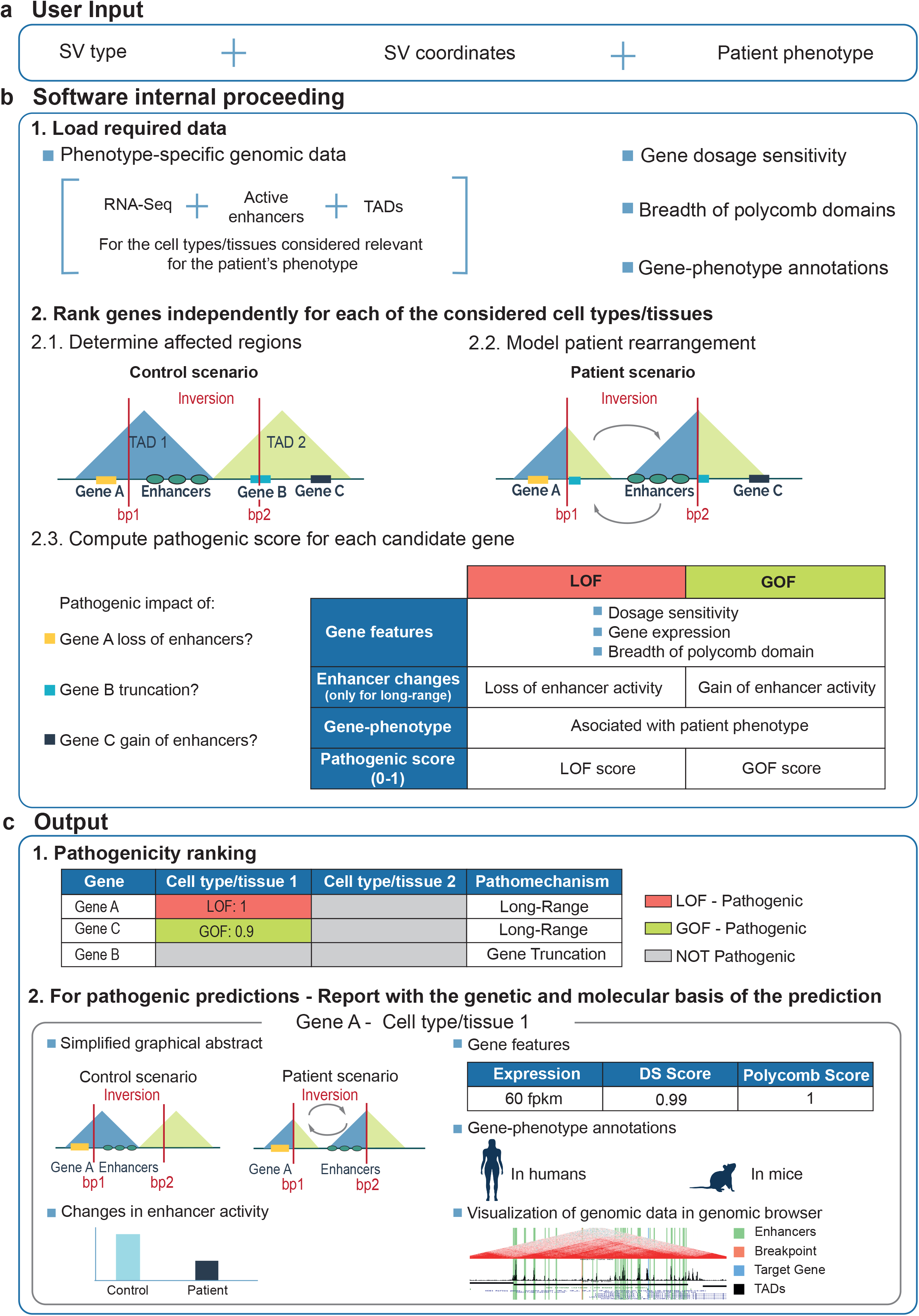
POSTRE Overview. **a,** *User Input:* For each SV identified in a patient, POSTRE requires (i) the type of SV (Deletion, Duplication, Inversion or Translocation), (ii) the genomic coordinates of the SV breakpoints at base-pair (bp) resolution and (iii) the patient’s phenotype. **b,** *Software internal proceeding:* cell types/tissues considered relevant for the patient’s phenotype are selected. For each of the selected cell types/tissues, specific genomic data (e.g. gene expression profiles, enhancer maps) are loaded to predict the SV pathogenicity. Each of the cell types/tissues considered as relevant for the patient’s phenotype is independently assessed. TADs are used as a proxy to determine the regulatory domains and genes potentially affected by the SV. To evaluate the pathogenicity of the candidate genes several features, such as dosage sensitivity (DS) or previous associations with the patient phenotype, are considered. In addition, for those candidate genes that are not directly disrupted by by the SV, long-range regulatory mechanisms resulting in either a gain (GOF) or loss (LOF) of gene expression are considered. **c,** *Output:* heatmap providing an overview of the predictions for each candidate gene and relevant cell type/tissue. Genes considered as pathogenic are highlighted in in red (Loss of Function, LOF) or green (Gain of Function, GOF). For genes predicted as pathogenic, a detailed report describing the genetic and molecular basis of the prediction is provided. The report includes a simplified graphical abstract of how the SV affects the candidate gene, details about enhancer changes in the regulatory domain of the candidate gene or a link to visualize relevant genomic data in a genome browser.

POSTRE requires three main inputs (Figure 1a): (i) the type of SV (i.e. deletion, duplication, inversion or translocation) (ii) the genomic coordinates of the SV breakpoints and (iii) the patient phenotype (i.e. limb, craniofacial/head&neck, cardiac or neurodevelopmental). Once the input data is submitted (Figure 1b), the cell types/tissues considered most relevant for the patient phenotype are selected for downstream analysis (Supplementary Data 1). Moreover, for each cell type/tissue, multiple developmental stages and/or *in vitro* differentiation time-points are typically analyzed. For instance, NCCs obtained at two different time-points through an *in vitro* differentiation protocol are used for the study of craniofacial abnormalities (Prescott et al., 2015; Laugsch et al., 2019), while two different developmental stages of the embryonic brain prefrontal cortex are analyzed for neurodevelopmental disorders (Markenscoff-Papadimitriou et al., 2020). POSTRE uses three main types of genomic information for each of the selected cell types/tissues: (i) gene expression profiles (based on RNA-Seq), (ii) active enhancer maps (based on ChIP-Seq) and (iii) TAD maps (based on Hi-C). Both gene expression profiles and active enhancer maps are specific for each cell type/tissue and developmental stage. However, due to the relative scarcity of Hi-C data, some of the TAD maps come from cellular contexts different than the ones relevant for each phenotype. This is justified by the general stability of TADs among different cell types (Dixon et al., 2012). In this regard, TAD maps generated in ESCs were successfully used to annotate regulatory domains disrupted by SVs associated with limb or craniofacial abnormalities (Lupiáñez et al., 2015; Laugsch et al., 2019). Once the relevant genomic data is loaded, the impact of the SV is independently evaluated for all the considered cell types/tissues. Firstly, the SV breakpoints are mapped to the genome attending to the location of TADs, which are used as a proxy to determine the regulatory domains affected by the genetic rearrangement. Subsequently, all the genes located in the disrupted TADs are considered as potentially affected by the SV (candidate genes) through either coding or long-range mechanisms. Then, all the candidate genes are ranked according to multiple features in order to estimate their likelihood of being involved in the patient disease. A summary of these features is presented below:

- **Gene-phenotype annotation:** genes previously associated with the patient phenotypic category (e.g. limb malformation) are considered as phenotypically relevant. This information is obtained from the Online Mendelian Inheritance in Man (OMIM) (Amberger et al., 2009) and the Mouse Genome Informatics (MGI) (Bult et al., 2019) databases.
- **Gene features:**

- *Dosage sensitivity (e.g. haploinsufficiency):* deviations from the normal dosage (i.e. number of copies), or expression levels, can be detrimental for some but not all genes. Hence, it is important to know whether the affected genes are dosage sensitive or not.
- *Breadth of polycomb domains in promoter regions:* genes whose promoters are covered by broad polycomb/H3K27me3 domains when inactive often correspond with major developmental genes implicated in congenital disorders (Rehimi et al., 2016; Shim et al., 2020). Moreover, this type of gene shows high enhancer responsiveness (Kraft et al., 2019; Pachano et al., 2021), which might be attributed to the presence of promoter CpG islands that prevent DNA methylation, facilitate enhancer-gene communication and, overall, provide a permissive chromatin environment.
- *Gene expression:* the expression status of the candidate genes is particularly relevant for LOF situations, as in those cases, the disease-causative genes must be expressed in the relevant cellular context to be involved in the patient condition.
- **Enhancer changes:** for the candidate genes that are not directly disrupted by the SV, long-range regulatory mechanisms are considered instead. Briefly, the enhancer activity associated to each candidate gene is estimated as the sum of the H3K27ac levels present at all the enhancers found within their TAD in the presence (patient) or absence (control) of the SV (Fulco et al., 2019). Then, differences in enhancer activity between the patient and control situations are computed for each candidate gene. For LOF, the candidate genes should lose enhancer activity (i.e. control enhancer activity > patient enhancer activity) through the deletion of cognate enhancers or enhancer disconnection pathomechanisms (Benko et al., 2009; Laugsch et al., 2019). For GOF, the candidate genes should gain enhancer activity (i.e. control enhancer activity < patient enhancer activity) through the duplication of cognate enhancers (Klopocki et al., 2008; Cox et al., 2011; Lohan et al., 2014) or enhancer adoption/hijacking pathomechanisms (Lettice et al., 2011).

Based on the previous criteria, each candidate gene will receive an overall pathogenic score (PS) between 0 and 1, with 1 meaning that it fulfills all the pathogenic criteria for a certain cell type/tissue (see Methods for more details). Then, these pathogenic scores are used to rank all the candidate genes across the considered cell types/tissues (Figure 1c). Moreover, those genes receiving a pathogenic score higher than a threshold (0.8) are highlighted as potentially pathogenic and a detailed report is provided for each of them. These reports contain a set of graphics, text and links to different external resources (e.g. UCSC genome browser with enhancer and TAD maps) to illustrate why a candidate gene is predicted as pathogenic in a specific cell type/tissue due to the SV (Supplementary Data 2).

Lastly, POSTRE can be executed with two alternative running modes: *Standard* or *High Specificity* (see Methods for details). Briefly, the *Standard* mode requires that, in order to receive high pathogenicity scores, the candidate genes display at least broad polycomb domains or high dosage sensitivity, while the *High-Specificity* mode requires that both criteria are fulfilled. In addition, POSTRE allows users to simultaneously analyze multiple SVs, each potentially associated with multiple phenotypes, in an automatic and sequential manner.

### POSTRE performance with experimentally validated SVs causing disease through long-range mechanisms

To illustrate POSTRE performance, we initially focused on a set of six patients with SVs causing congenital abnormalities (limb or craniofacial) compatible with POSTRE’s analysis pipeline (Table 1a; Supplementary Data 3). Importantly, each of these SVs only alters pathogenically the expression level of a single gene, through experimentally validated long-range pathomechanisms (Lohan et al., 2014; Lupiáñez et al., 2015; Franke et al., 2016; Laugsch et al., 2019; Long et al., 2020). A summary of POSTRE’s results obtained for these six patients is shown in Table 1b-c.

**Table 1.** Overview of POSTRE analyses for SVs causing congenital abnormalities through experimentally validated long-range mechanisms.

**a) Patients description**
| Patient Nr | SV Type | Causative Gene | Pathological mechanism | Phenotype |
| --- | --- | --- | --- | --- |
| 1 | Inversion | *TFAP2A* | Downregulation by enhancer disconnection | Head&Neck |
| 2 | Deletion | *SOX9* | Downregulation by enhancer deletion | Head&Neck |
| 3 | Duplication | *KCNJ2* | Upregulation by enhancer adoption | Limbs |
| 4 | Deletion | *PAX3* | Upregulation by enhancer adoption | Limbs |
| 5 | Duplication | *IHH* | Upregulation by enhancer adoption | Limbs |
| 6 | Duplication | *SHH* | Upregulation by enhancer duplication | Limbs |

**b) Information candidate genes**
| Patient Nr | N candidate genes based on TADs (causative + non-causative) |
| --- | --- |
| 1 | 6 |
| 2 | 1 |
| 3 | 5 |
| 4 | 5 |
| 5 | 46 |
| 6 | 7 |
| Total: | 70 |

**c) Summary of POSTRE results for the six patients**
| For causative genes (N=6) |  |
| --- | --- |
| N causative genes identified by POSTRE as causative | % causative genes identified correctly |
| 5 | 83% (Sensitivity) |

| For non-causative genes (N = 64) |  |
| --- | --- |
| N non-causative genes identified by POSTRE as non-causative | % non-causative genes identified correctly |
| 64 | 100% (Specificity) |

After mapping the breakpoints of the six SVs with respect to TADs in the relevant cell types/tissues, a total of 70 candidate genes were identified (Table 1b). Remarkably, for five out of six patients, POSTRE successfully predicted the single causative gene whose expression is affected by each SV as well as the implicated long-range pathomechanism (Table 1c). Overall, POSTRE achieved a sensitivity of 83% and a specificity of 100% (i.e. no false positive genes predicted) at the gene level for this limited, albeit relevant, patient group (Table 1c). For example, for a patient with craniofacial abnormalities carrying a heterozygous inversion in Chr6 (i.e. *TFAP2A* patient; Table 1a; patient Nr 1 in Supplementary Data 3), POSTRE predicted the physical disconnection between *TFAP2A* and some of its cognate enhancers in NCC (i.e. enhancer disconnection) (Figure 2a-d). POSTRE’s report indicates that this enhancer disconnection can lead to the loss of *TFAP2A* expression in NCC and the emergence of craniofacial defects, in agreement with the experimentally validated pathomechanism (Laugsch et al., 2019). POSTRE assigned *TFAP2A* a high pathogenic score because (i) it is highly expressed in NCC, (ii) it loses enhancer activity in NCC due to the inversion, (iii) it is a dosage sensitive gene, (iv) its promoter displays a broad polycomb domain when it is inactive and (v) it has been previously associated with craniofacial (head&neck) abnormalities in OMIM and MGI (Figure 2a-d; Supplementary Figure 1a-f; Supplementary Data 2). Similarly, for a patient with limb abnormalities carrying a heterozygous deletion in Chr2 (i.e. *PAX3* patient; Table 1a; patient Nr 4 in Supplementary Data 3), POSTRE successfully predicted a pathogenic gain of *PAX3* expression in the limb due to an enhancer adoption mechanism (Figure 3a-d). It is worth mentioning that, although the deletion eliminates one of the *EPHA4* alleles, POSTRE did not predict this gene as pathogenic. This is in perfect agreement with the experimental data (Lupiáñez et al., 2015) and highlights the relevance of considering long-range mechanisms even when protein-coding genes are directly affected by SVs. POSTRE assigned *PAX3* a high pathogenicity score because (i) it gains enhancer activity, (ii) it is a dosage sensitive gene, (iii) its promoter displays a broad polycomb domain when it is inactive and (iv) it has been previously associated with limb abnormalities in OMIM and MGI (Figure 3a-d; Supplementary Figure 2a-e).

**Figure 2.**
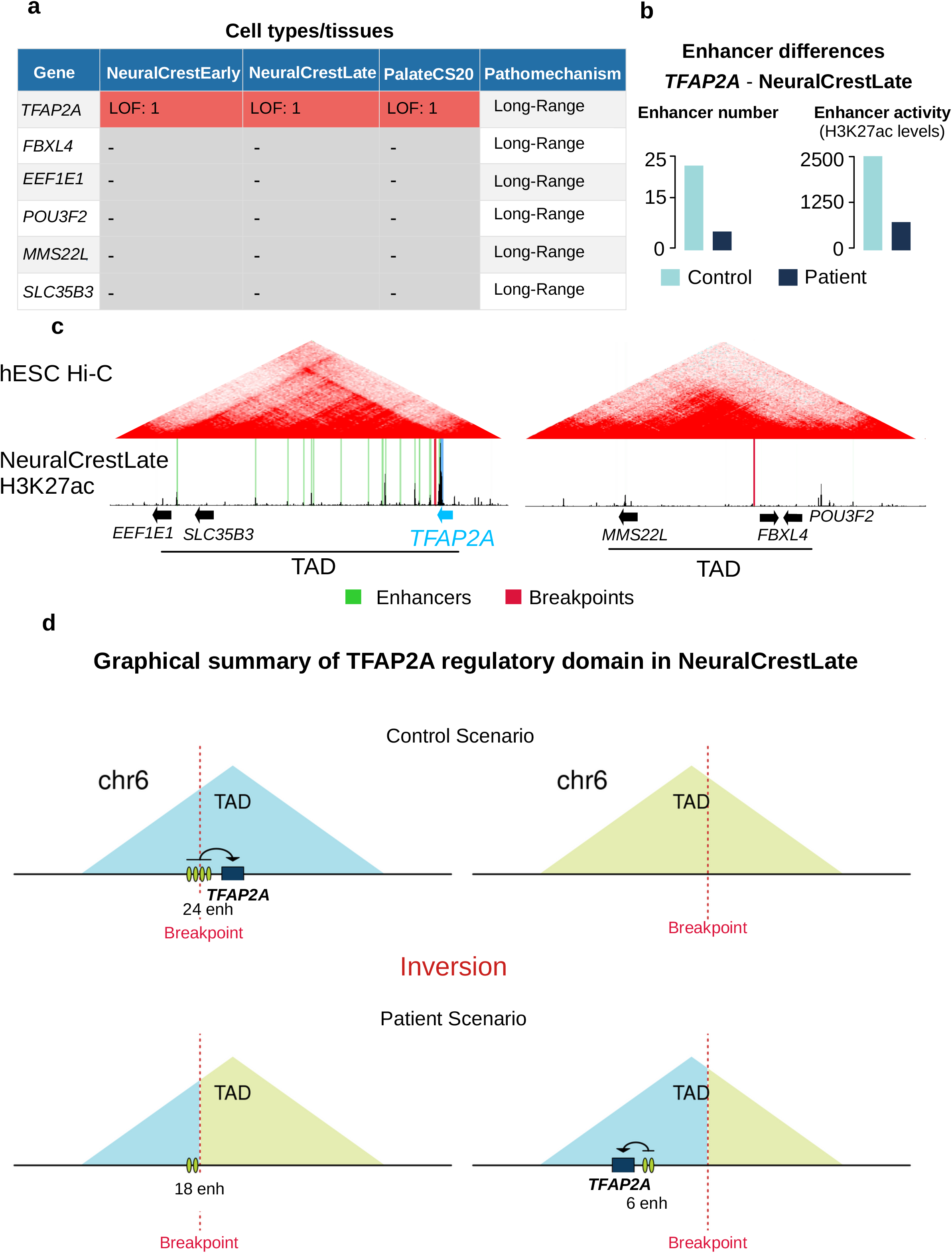
POSTRE results for the *TFAP2A* patient. **a**, An inversion in Chr6 was previously shown to cause craniofacial abnormalities through the disconnection between *TFAP2A* and its Neural Crest Cell (NCC) cognate enhancers, resulting in the haploinsufficient expression of *TFAP2A* in NCC (Laugsch et al., 2019). The inversion was analyzed by POSTRE and, among the candidate genes, *TFAP2A* was the only one considered as pathogenic in two human NCC *in vitro* differentiation stages (NeuralCrestEarly and NeuralCrestLate) and the embryonic palate from Carnegie stage 20 (Palate CS20) (See Supplementary Data 1 for more details). **b**, Enhancer activity (H3K27ac levels; see Methods) and number of enhancers associated with *TFAPA2A* in NCC (NeuralCrestLate) are shown in the absence (Control) or presence (Patient) of the inversion. **c**, Genome browser view of the two TADs affected by the inversion. The inversion breakpoints are highlighted in red and the *TFAP2A* cognate enhancers in NeuralCrestLate in green. **d**, Graphical abstract illustrating the changes in the regulatory landscape of *TFAP2A* due to the inversion in NeuralCrestLate. The green ovals represent enhancers (enh).

**Figure 3.**
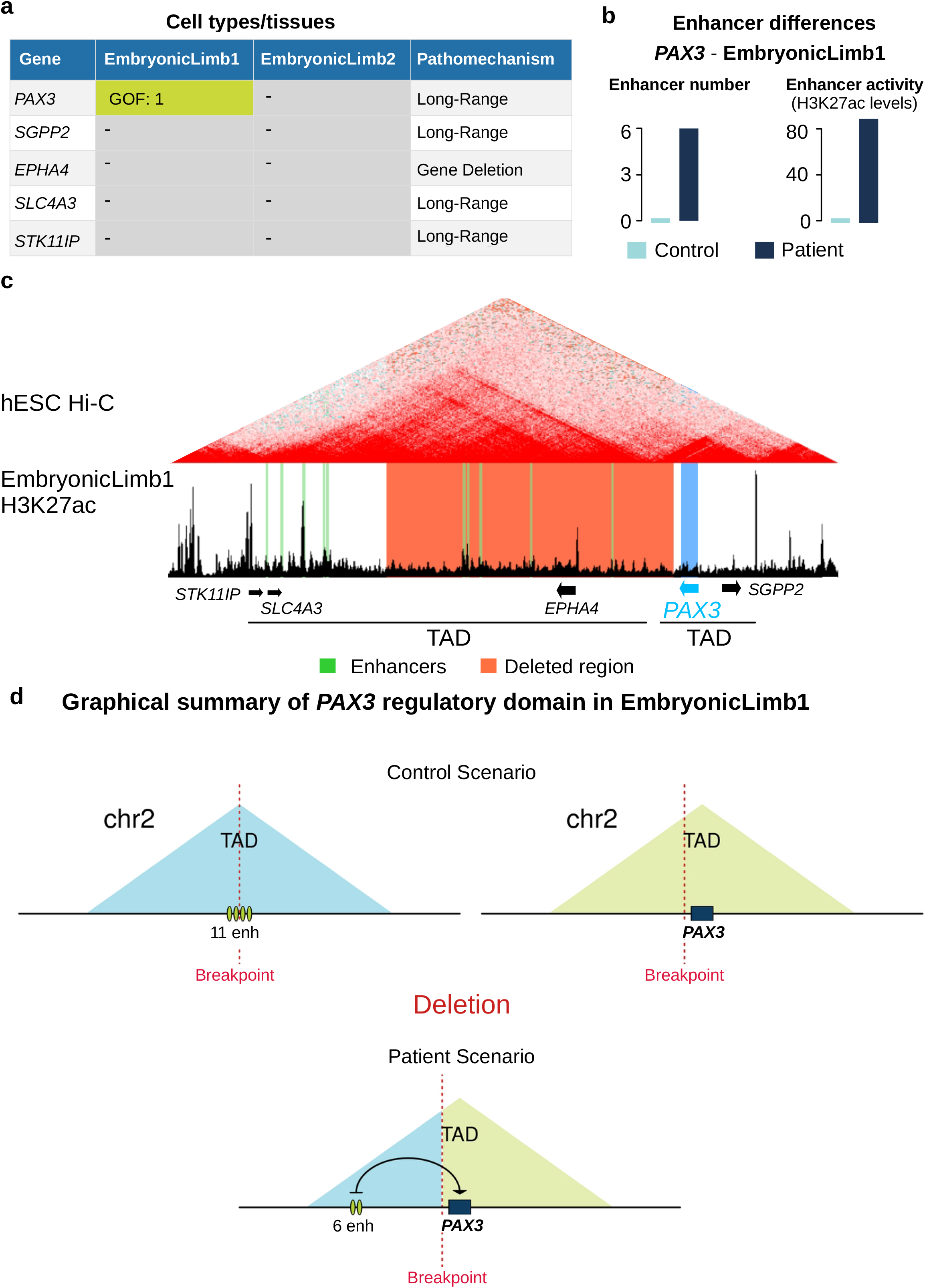
POSTRE results for the *PAX3* patient. **a,** A deletion in Chr2 was previously shown to cause limb abnormalities through an enhancer adoption mechanism leading to *PAX3* ectopic expression in the developing limb bud (Lupiáñez et al., 2015). The deletion was analyzed by POSTRE and, among the candidate genes, *PAX3* was the only one considered as pathogenic in two human limb bud developmental stages (EmbryonicLimb1, EmbryonicLimb2) (See Supplementary Data 1 for more details). **b**, Enhancer activity (H3K27ac levels) and number of enhancers associated with *PAX3* in embryonic limb buds (EmbryonicLimb1) are shown in the absence (Control) or presence (Patient) of the deletion. **c**, Genome browser view of the two TADs affected by the deletion. The deletion is highlighted in orange and the limb bud active enhancers in EmbryonicLimb1 in green. **d**, Graphical abstract illustrating the changes in the regulatory landscape of *PAX3* in EmbryonicLimb1 due to the deletion. The green ovals represent enhancers (enh).

For one of the six analyzed patients POSTRE did not predict the causative gene due to limitations in the available genomic data (Patient Nr 6 in Table 1a and Supplementary Data 3). Briefly, previous work showed that in this patient a duplication in Chr7 spanning the ZRS enhancer lead to abnormally high *SHH* expression levels in the embryonic limb (Lohan et al., 2014). Notably, the ZRS enhancer is active and controls *SHH* expression in a rather limited number (<4%) of cells (i.e. zone of polarizing activity (ZPA)) within the developing limb bud (Vandermeer et al., 2014). In contrast, the genomic data used by POSTRE was generated from bulk human limb buds rather than isolated ZPA cells. As a result, it was not possible to detect the ZRS enhancer, which precluded the prediction of *SHH* as the relevant gene in this patient.

### POSTRE performance with SVs previously predicted to cause human congenital abnormalities through long-range mechanisms

Next, we focused on a set of 38 patients in whom SVs were previously predicted to cause congenital abnormalities through long-range mechanisms that, nevertheless, have not been experimentally validated (Table 2; Supplementary Data 4). In a subset of these patients, the associated SVs recurrently disrupt the same regulatory domains and potentially alter the expression of disease-relevant genes (e.g. *SATB2*, *MEF2C*, *FOXG1*, *DLX5&6*) (D’haene and Vergult, 2021). For other patients, the impact of the associated SVs on the expression of the putative causative genes (e.g. *SLC2A1*, *BCL11B*) was assessed, but not in the appropriate cell type/tissue for the investigated diseases (e.g. analysis in patient blood cells for a neurodevelopmental disorder) (Redin et al., 2017; Lessel et al., 2018; D’haene and Vergult, 2021).

**Table 2.**
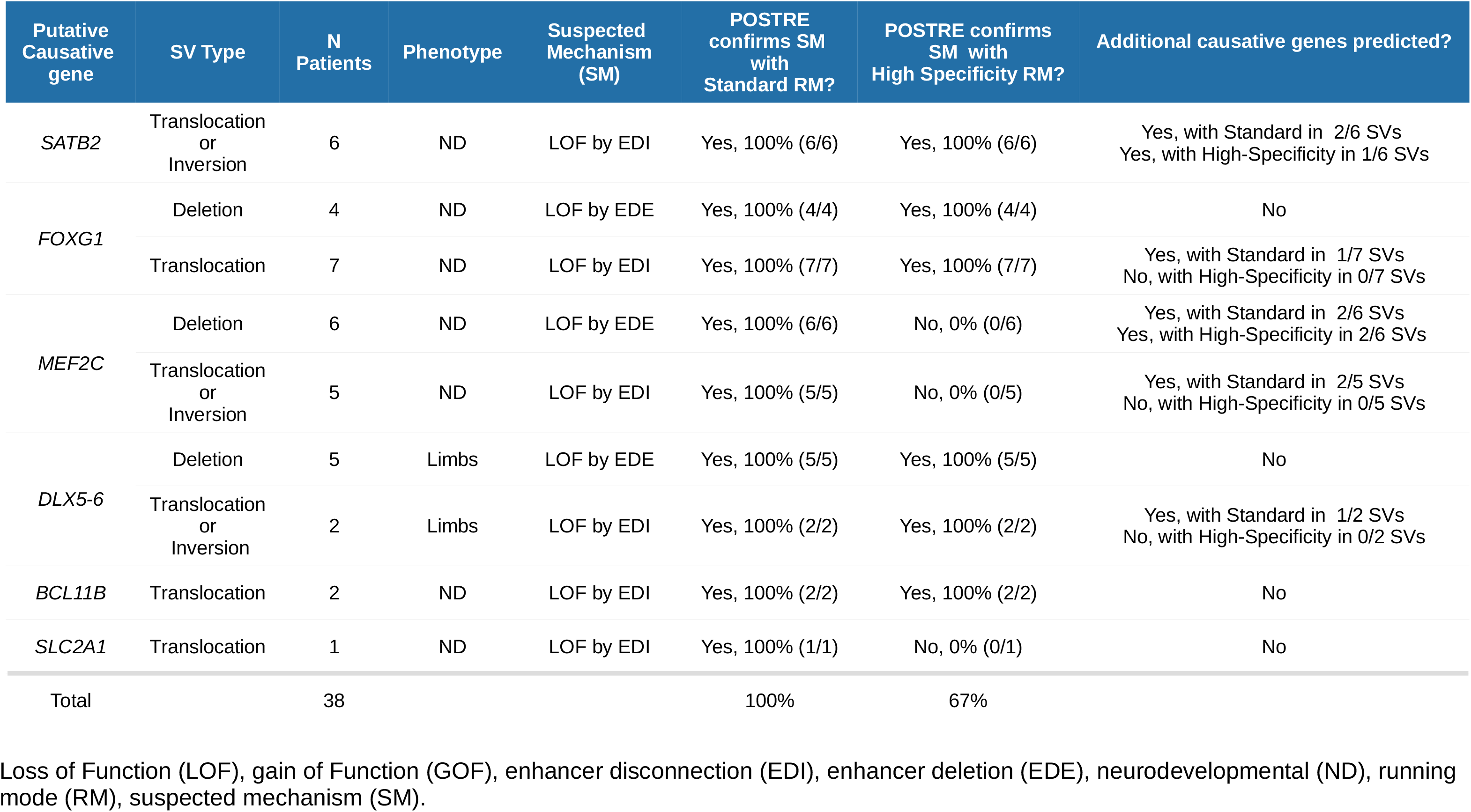
Overview of POSTRE analyses for SVs previously predicted to cause congenital abnormalities through long-range mechanisms.

When analyzing these 38 patients with POSTRE using the *Standard* mode, the previously proposed causative genes and long-range pathomechanisms were successfully predicted in 100% of the cases (Table 2). This was true regardless of whether the SV breakpoints were close or distally located with respect to the target genes, as depicted for *SATB2* (Supplementary Figure 3a-c). The sensitivity dropped to 67% when using the *High specificity* mode. The reason for this reduction is that two of the causative genes, *MEF2C* and *SLC2A1*, do not display broad polycomb domains when they are not active, one of the gene features required by the *High specificity* mode. Among the analyzed patients, it is worth highlighting one of them (DECIPHER ID: 260836; *FOXG1* patient Nr 8 in Supplementary Data 4) carrying a deletion located close to *FOXG1* and showing neurodevelopmental deffects. There has been some discrepancy regarding the mechanism whereby this deletion might affect *FOXG1* expression. It was originally proposed that the deletion could eliminate a TAD boundary and cause *FOXG1* overexpression through an enhancer adoption mechanism (Ibn-Salem et al., 2014). However, subsequent work indicated that this deletion could lead to *FOXG1* downregulation by eliminating some of its cognate brain enhancers (Mehrjouy et al., 2018), which is in agreement with *FOXG1* haploinsufficiency causing brain congenital abnormalities (e.g. Rett Syndrome) (Kumakura et al., 2014). Importantly, POSTRE also predicted that this deletion could cause a loss of *FOXG1* expression in the brain prefrontal cortex through the deletion of relevant enhancers (Supplementary Figure 4a-g).

In addition to the accurate identification of the previously proposed causative genes, POSTRE predicted a few additional genes for some of the analyzed SVs (Table 2, Figure 4). For example, for one of the deletions predicted to cause the loss of *MEF2C* expression in the prefrontal cortex through a long-range mechanism (*MEF2C* Nr 11 in Supplementary Data 4), POSTRE also identified *NR2F1* as a potentially pathogenic gene (Supplementary Figure 5a-h). In this case, POSTRE predicted a coding LOF mechanism, as the deletion eliminates one of the *NR2F1* alleles. Both *MEF2C* and *NR2F1* have been previously associated with neurodevelopmental deffects (Tocco et al., 2021; Zhang and Zhao, 2022). Therefore, the deletion could cause the loss of both *NR2F1* and *MEF2C* function through distinct mechanisms, which in turn might define the phenotypic spectrum of this patient. This illustrates how direct and long-range effects could be simultaneously elicited by a single SV (Ibn-Salem et al., 2014; Middelkamp et al., 2019).

**Figure 4.**
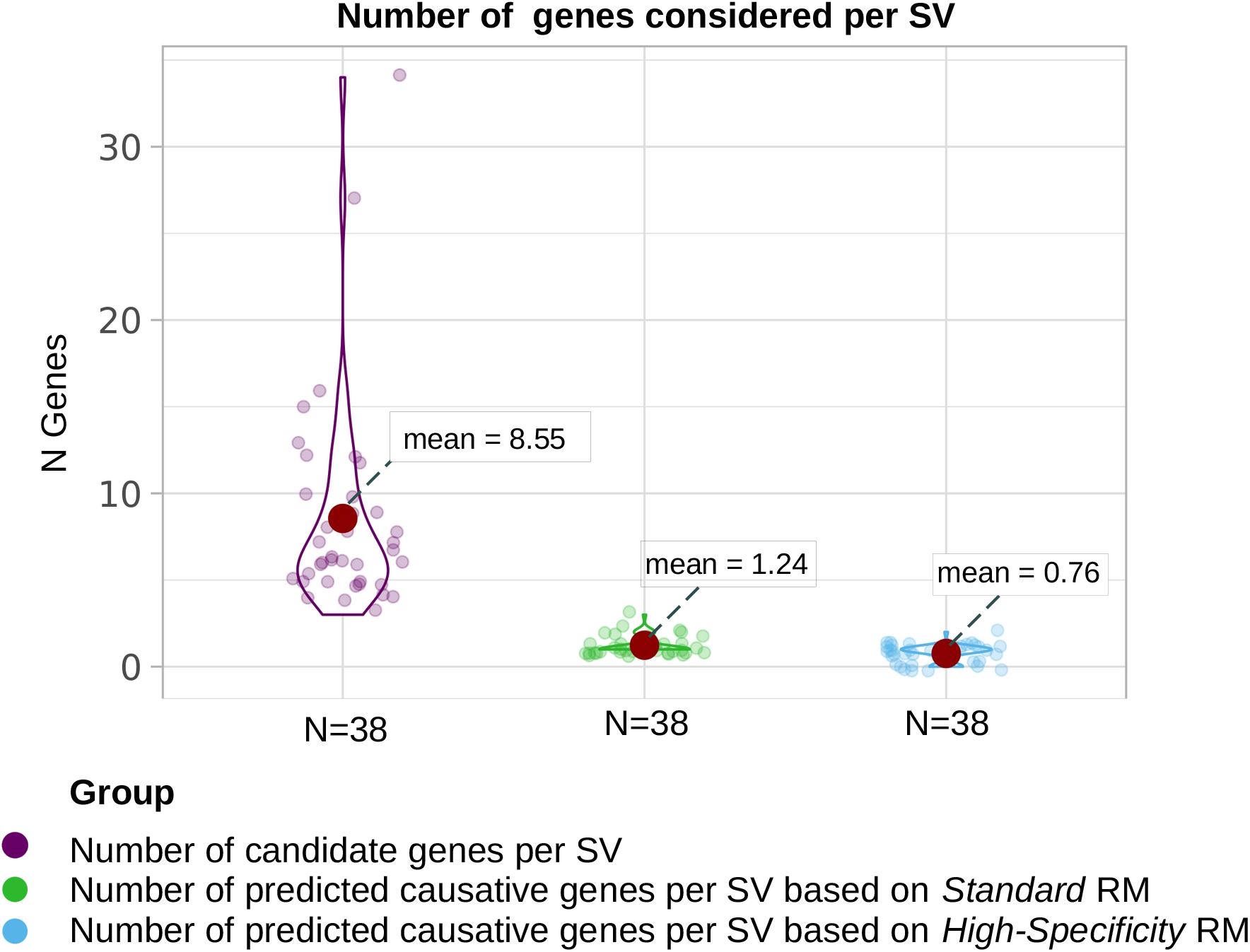
Overall number of candidate and pathogenic genes identified by POSTRE when analyzing SVs previously predicted to cause human congenital abnormalities. 38 SVs previously reported to cause congenital abnormalities through long-range pathomechanisms (Table 2) were analyzed by POSTRE. The total and average number of candidate genes located within the affected TADs is shown for each SV (purple). The total and average number of genes predicted as causative using either the *Standard* or *High Specificity* running modes (RM) are shown in green and blue, respectively. Genes defined as causative displayed pathogenic scores >0.8.

These examples (*FOXG1* Nr 8 and *MEF2C* Nr 11) highlight that, in contrast to previous approaches (Ibn-Salem et al., 2014), it is important to consider all possible mechanisms (i.e. coding LOF, coding GOF, long-range LOF, long-range GOF) when evaluating the pathogenic impact of SVs, as this might provide a better understanding of the etiology and phenotypic variability of human disease.

### Assessment of Type-1 error rate (false positives)

Having shown that POSTRE can effectively distinguish between pathogenic and non-pathogenic genes when applied to previously reported detrimental SVs (Table 1), we then wanted to assess POSTRE capacity to discriminate between non-pathogenic and pathogenic SVs. This is particularly important considering the abundance of structural variation in humans, with a typical genome carrying around 10.000 SVs (Lappalainen et al., 2019; Collins et al., 2020). For this purpose, SVs from healthy individuals available in gnomAD (Karczewski et al., 2020) were selected, as the majority of these variants are expected to be non-pathogenic. Firstly, we randomly selected 10.000 SVs and analyzed them using the four different phenotypic categories currently handled by POSTRE (i.e. craniofacial/head&neck, neurodevelopmental, limb and cardiovascular). Chiefly, POSTRE did not predict any pathogenic effect for most of the SVs (97.8% non-pathogenic for the *Standard* mode; 99.2% non-pathogenic for the *High Specificity* mode) (Table 3a). In general, pathogenic SVs tend to be larger and, consequently, affect a higher number of genes than non-pathogenic ones (Rodriguez-Revenga et al., 2007; Ibn-Salem et al., 2014). This size difference is readily observed between previously reported pathogenic SVs analyzed with POSTRE (Table 1 and 2) and those selected from healthy individuals (Table 3b). To evaluate whether these size differences could fully explain the low % of pathogenic predictions among SVs from healthy individuals, we also tested POSTRE performance after selecting the largest SVs present among healthy individuals. Briefly, we selected SVs >75 Kb (size of the smallest SV analyzed in Table 1a) obtaining 2980 SVs, as well as the top 500 and top 100 largest SVs from gnomAD. Importantly, POSTRE still reported most of these SVs as non-pathogenic, even for the top 100 largest SVs (98% for *High-Specificity* mode; 86% for *Standard* mode) (Table 3a). Altogether, these results show that POSTRE controls well the Type-1 errors (false positives).

**Table 3.** Analysis of non-pathogenic (control) SVs with POSTRE.

**a) POSTRE prediction results**
|  | % of SVs identified as Not Pathogenic |  |
| --- | --- | --- |
|  | Standard RM | High Specificity RM |
| **Subset 1 (10K controls)** | 97.8% | 99.2% |
| **Subset 2 (2980 controls)** | 95.0% | 98.7% |
| **Subset 3 (500 controls)** | 92.2% | 97.9% |
| **Subset 4 (100 controls)** | 86.0% | 98.0% |

**b) Analysis of the size distribution for the different groups of SVs in base pairs**
|  | 1st Quartile | Median | 3rd Quartile |
| --- | --- | --- | --- |
| **All gnomAD controls (>100K)** | 107 | 547 | 3566 |
| **Subset 1 (10K controls)** | 106 | 560 | 3538 |
| **Subset 2 (2980 controls)** | 102468 | 148862 | 268102 |
| **Subset 3 (500 controls)** | 426885 | 543023 | 822252 |
| **Subset 4 (100 controls)** | 1093246 | 1377514 | 2577491 |
| **Pathogenic SVs Table 1 & 2** | 404592 | 887812 | 2409324 |

### Database of POSTRE predictions for patients with congenital abnormalities carrying SVs

POSTRE can be used to analyze large cohorts of patient SVs described in the literature and/or deposited in databases (Firth et al., 2009; Lappalainen et al., 2013; Redin et al., 2017; Landrum et al., 2018; Middelkamp et al., 2019), many of which are currently considered as variants of uncertain significance (VUS) (Federici and Soddu, 2020). The systematic analysis of large SV cohorts might (i) identify relevant disease loci based on recurrence, (ii) predict novel pathological mechanisms for already known disease-relevant genes and (iii) provide testable hypothesis regarding the pathogenicity of VUS. To illustrate this, we analyzed a set of 270 patients described in (Redin et al., 2017; Middelkamp et al., 2019) and displaying phenotypes compatible with POSTRE. POSTRE predicted pathogenic events in 53% and 19,4% of the patients with the *Standard* and *High-Specificity* modes, respectively (Figure 5a), which is considerably higher than for the healthy controls described in Table 3. Moreover, POSTRE did not simply identify SVs with pathogenic potential, but also predicted the cellular context in which the SVs might elicit their pathogenic effects as well as their potential mechanism of action. In this regard, when considering all the predicted pathological events with the *Standard* running mode, 49% corresponded to long-range regulatory effects (Figure 5b), highlighting the potential prevalence of this type of pathomechanism. On the other hand, when considering POSTRE predictions (*Standard* running mode) for all 270 patient SVs, the average number of candidate genes was 31.68, while the average number of causative genes was 1.71 (Figure 5c). The average number of candidate genes was clearly higher than the one described for patients analyzed in previous sections (Figure 4), probably because the SVs in these 270 patients also tend to be larger (Supplementary Figure 6). However, the number of predicted causative genes was rather similar between these two patient cohorts (1.71 vs 1.24; Figure 5c and Figure 4). This suggests that POSTRE can facilitate the prioritization of disease-relevant genes even when many candidate genes are potentially affected by large SVs. Nevertheless, approximately half of the patient SVs were not predicted as pathogenic, which highlights the need to better understand the non-coding genome and to incorporate genomic data from additional human cell types in order to reveal the full pathogenic spectrum of structural variation.

**Figure 5.**
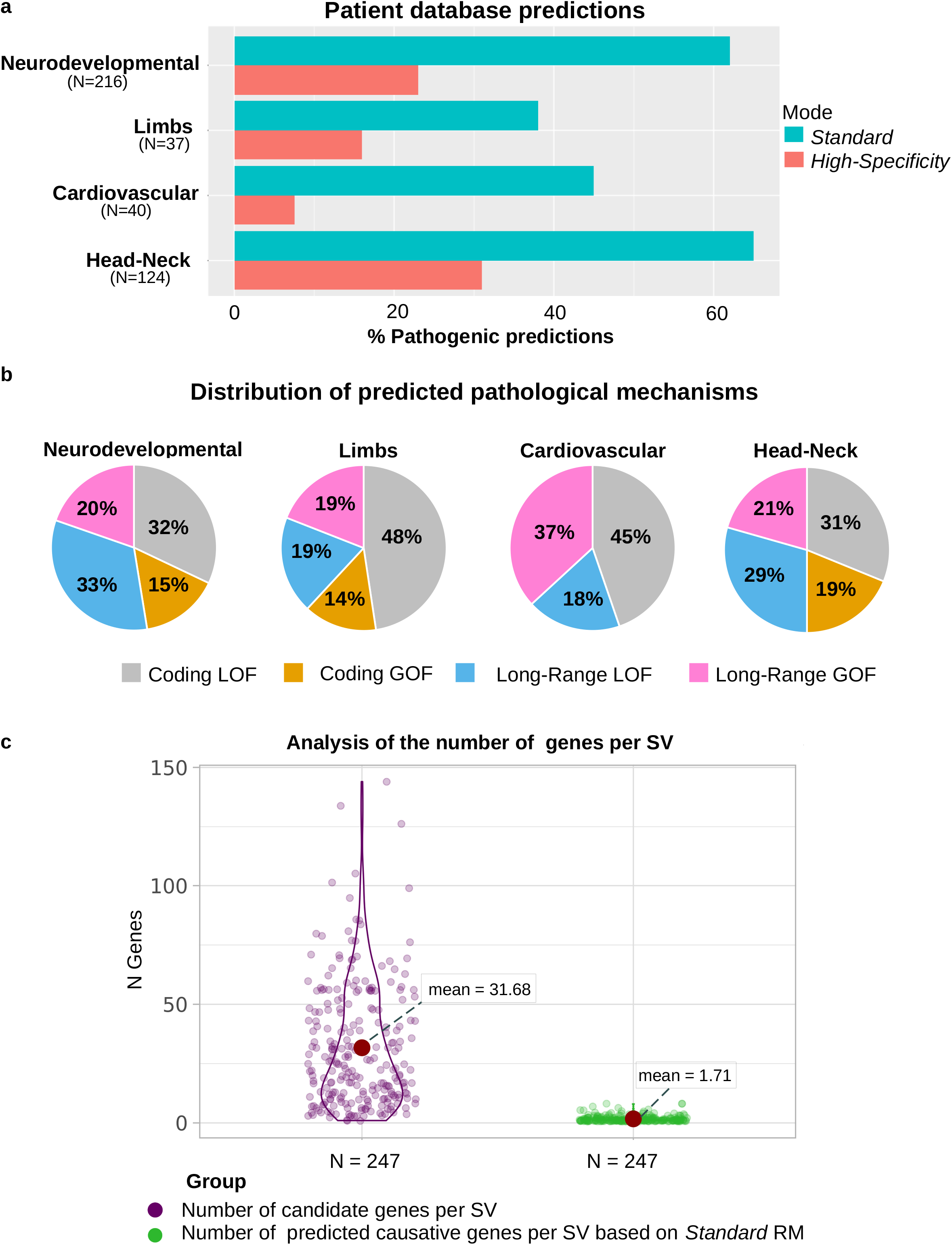
POSTRE analyses for a cohort of 270 patient SVs. **a**, Percentage of pathogenic SV found by POSTRE for the indicated phenotypes using the *Standard* or *High-Specificty* modes. N=total number of SVs associated with each of the indicated phenotypes. **b**, Pie charts showing the relative abundance of the main pathological mechanisms (i.e. coding GOF, coding LOF, long-range GOF, long-range LOF) among the SVs predicted as pathogenic by POSTRE for each of the indicated phenotypes using the *Standard* mode. **c**, Violin plots showing the number of candidate (purple, left) and causative (green, right) genes identified by POSTRE (*Standard* running mode) for each of the analyzed SVs predicted to be pathogenic. Each dot corresponds to a SV.

For more than half of the 270 analyzed patients, POSTRE predicted a pathogenic SV as well as an underlying pathomechanism, which can improve the current understanding of these disorders and facilitate the design of experimental strategies to uncover their molecular etiology (Sánchez-Gaya et al., 2020). All these predictions are available through POSTRE’s *Explore Previous SVs* section, enabling users to browse them by gene, phenotype, pathological mechanism and cell type/tissue. To illustrate the usefulness of these results, we now describe POSTRE predictions for a couple of the investigated patients:

*Patient UTR8 (dbVar):* patient with Pierre Robin sequence (PRS), a syndrome associated with craniofacial abnormalities. This patient carries a translocation (UTR8 in dbVar, 268030 in ClinVar) considered as a VUS in ClinVar (Figure 6a). POSTRE predicted that this translocation could cause a pathogenic loss of *SOX9* expression in NCC through an enhancer disconnection mechanism (Figure 6b-d). This prediction is supported by previous work (Long et al., 2020), where some of the NCC enhancers that are disconnected from *SOX9* due to the translocation were deleted in hESC using CRISPR/Cas9 technology. Notably, the deletion of those enhancers significantly and specifically reduced *SOX9* expression upon differentiation of hESC into NCC, supporting that *SOX9* LOF through long-range mechanisms could lead to PRS.

**Figure 6.**
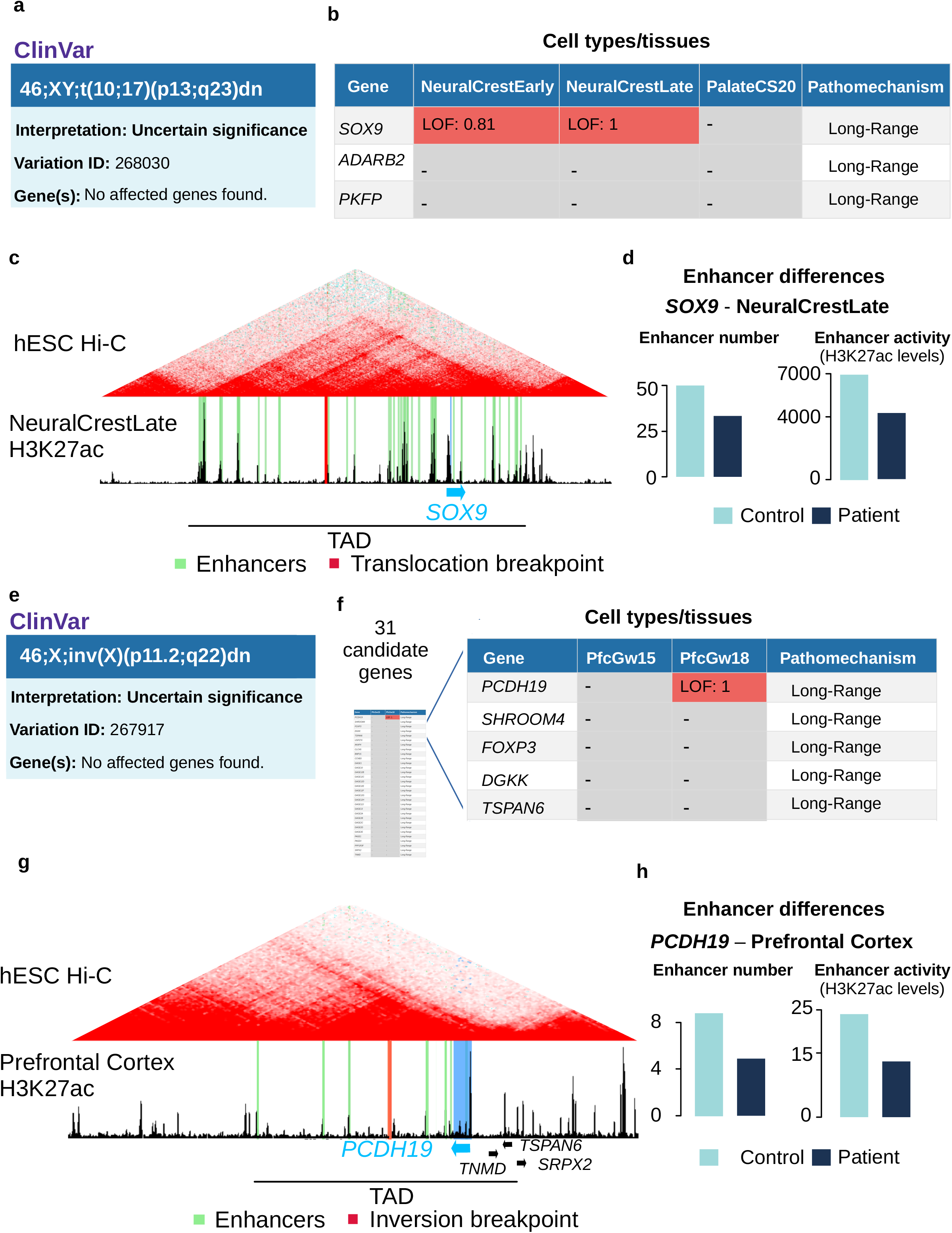
Long-Range LOF predictions for UTR8 and DGAP270 patients. **a,** Patient UTR8 displays craniofacial abnormalities (Pierre Robin Sequence) and carries a translocation with breakpoints in Chr10 and Chr17, which is currently classified as a VUS in ClinVar. **b,** The UTR8 translocation was analyzed by POSTRE and, among the candidate genes, *SOX9* was the only one considered as pathogenic in two NCC *in vitro* differentiation stages (NeuralCrestEarly, NeuralCrestLate) (See Supplementary Data 1 for more details). **c**, Genome browser view of one of the TADs affected by the UTR8 translocation, in which *SOX9* is located. The translocation breakpoint is highlighted in red and the NCC active enhancers in NeuralCrestLate in green. **d**, Enhancer activity (H3K27ac levels) and number of enhancers associated with *SOX9* in NCC (NeuralCrestLate) are shown in the absence (Control) or presence (Patient) of the UTR8 translocation. **e,** Patient DGAP270 displays neurodevelopmental defects and carries an inversion in ChrX, which is currently classified as a VUS in ClinVar. **f,** The DGAP270 inversion was analyzed by POSTRE and, among the candidate genes, *PCDH19* was the only one considered as pathogenic in the embryonic prefrontal cortex gestation week 18 (PfcGw18) (See Supplementary Data 1 for more details). **g**, Genome browser view of one of the TADs affected by the DGAP270 inversion, in which *PCDH19* is located. The inversion breakpoint is highlighted in red and the prefrontal cortex active enhancers in PfcGw18 in green. **h**, Enhancer activity (H3K27ac levels) and number of enhancers associated with *PCDH19* in the prefrontal cortex (PfcGw18) are shown in the absence (Control) or presence (Patient) of the DGAP270 inversion.

*Patients U152 and DGAP270 (dbVAR):* patients with neurodevelopmental defects carrying a deletion (U152, dbVar) or an inversion (DGAP270, dbVar) in ChrX. In both cases, POSTRE predicted LOF for *PCDH19* in the brain prefrontal cortex through either coding (U152) or long-range regulatory mechanisms (DGAP270). PCDH19 is associated with neurodevelopmental disorders, especially with epilepsy and intellectual disability (Smith et al., 2018; Symonds et al., 2019; Samanta, 2020). In the case of DGAP270, the associated inversion was listed as a VUS in ClinVar (ID: 267917) (Figure 6e). Notably, POSTRE predicted that the inversion could cause a loss of PCDH19 expression in the brain prefrontal cortex through an enhancer disconnection mechanism (Figure 6f-h).

### POSTRE benchmarking

Computational tools developed to analyze SVs can be broadly divided in two main, non-mutually exclusive, categories. On the one hand, there are “annotation” tools that provide different layers of descriptive information for the genomic regions affected by the SVs (e.g. VEP (McLaren et al., 2016), ClinTAD (Spector and Wiita, 2019), TADA (Hertzberg et al., 2022), TADeus2 (Poszewiecka et al., 2022) or POSTRE). The type of provided information differs among tools, but can include: the genes directly affected or located close to the SV, the recurrence of SVs in the same locus for additional patients, the presence of *cis*-regulatory elements, etc. Although “annotation” tools can help characterizing SVs of interest, they require the user to integrate the provided information and to predict a plausible pathomechanism accordingly. On the other hand, there are “prediction” tools that can rank and prioritize SVs based on their pathogenicity, although only a few of them consider both coding and long-range pathomechanisms (e.g. POSTRE, TADA (Hertzberg et al., 2022), TADeus2 (Poszewiecka et al., 2022), SVScore (Ganel et al., 2017)). A more detailed comparison between POSTRE and other available tools is presented in Table 4 and briefly discussed below.

**Table 4.** POSTRE comparison with other SV analysis tools.

|  | POSTRE | StrVCTVRE | VEP | ClinTAD | SVScore | TADA | TADeus2 |
| --- | --- | --- | --- | --- | --- | --- | --- |
| Types of SVs | Del, Dup, Inv, Trans | Del, Dup | Del, Dup | Del, Dup | Del, Dup, Inv, Trans | Del,Dup | Del,Dup, Inv,Trans |
| Descriptive-Annoation Tool | + | - | + | + | - | + | + |
| Prediction of pathogenic coding effects | + | + | + | - | + | + | + |
| Prediction of pathogenic long-range effects | + | - | - | - | +/- | +/- | +/- |
| Considers patient phenotype | + | - | - | + | - | - | - |
| Applicable to all diseases | - | + | + | + | + | + | + |
| GUI User-friendly | + | + | + | + | - | - | + |
| Accepts Multiple SVs | + | - | + | + | + | + | - |
| Cellular context dependent predictions | + | - | - | - | - | - | - |
| Cellular context dependent report explaining predictions | + | - | - | - | - | - | - |

Deletion (Del), duplication (Dup), inversion (Inv), translocation (Trans), + (Yes), - (No), +/- (Limited), graphical user interface (GUI)

#### Pathomechanisms

Several tools, including POSTRE, are able to predict whether SVs have direct effects on protein coding genes (e.g. changes in gene dosage or gene disruption), although methodological differences may exist among each of them. For instance, StrVCTVRE (Sharo et al., 2022) quantifies the percentage of exons affected by the SVs, while POSTRE simply considers whether the gene sequence is affected or not. On the other hand, both StrVCTVRE (Sharo et al., 2022) and POSTRE require that strong candidate genes display dosage sensitivity.

Despite the accumulating evidences indicating that SVs can frequently cause disease through long-range mechanisms, most computational tools can not make this type of predictions (e.g. StrVCTVRE (Sharo et al., 2022)). Furthermore, SVs can cause human disease through long-range regulatory mechanisms even when they directly affect a protein coding gene (Figure 3a) (Lupiáñez et al., 2015). Some available tools can annotate or rank SVs based on the presence of enhancers, or other cis-regulatory elements, within the genomic region disrupted by a SV (SVScore (Ganel et al., 2017), TADA (Hertzberg et al., 2020), TADeus2 (Poszewiecka et al., 2022)). However, these tools present some important limitations when predicting long-range pathological mechanisms. For instance, since TADA was developed by training a classifier with CNVs acting preferentially through coding pathomechanisms, the authors recommend the usage of their ranking model for coding rather than non-coding pathomechanisms (Hertzberg et al., 2020). Both SVScore (Ganel et al., 2017) and TADeus2 (Poszewiecka et al., 2022) consider long-range LOF (e.g. enhancer disconnection) but not long-range GOF (i.e. enhancer adoption) pathomechanisms. Therefore, POSTRE is rather unique, as it can identify pathogenic SVs causing gene expression changes through all major types of long-range mechanisms (i.e. enhancer adoption, enhancer disconnection, enhancer deletion, enhancer duplication).

#### Cellular context

For any given disease, the patients phenotype typically reflect the cell type/tissue in which the SVs have a detrimental effect. Considering the cellular context in which a SV might manifest its pathogenicity is particularly relevant in the case of long-range pathomechanisms, as enhancers display higher tissue specificity than genes. However, with a few exceptions (Ibn-Salem et al., 2014; Middelkamp et al., 2019), most available tools do not consider the patient phenotype when estimating the pathogenicity of SVs. For instance, as described in previous sections (Table1; Figure 3), POSTRE successfully predicted the pathogenicity of a deletion present in a patient with limb abnormalities that increases *PAX3* expression in the limb through an enhancer adoption mechanism (Lupiáñez et al., 2015). When this deletion was analyzed with TADeus2, *PAX3* but also additional genes, such as *EPHA4*, were predicted as disease-relevant. However, previous work conclusively showed that *PAX3* was the causative gene in this patient, with *EPHA4* not being involved in the limb abnormalities (Lupiáñez et al., 2015). In comparison to POSTRE, TADeus2 shows lower specificity probably because it does not consider the patient phenotype or uses tissue-specific genomic data. POSTRE takes into account the patient phenotype in order to select the genomic data and cellular context/s in which the impact of the SV should be modeled, as well as to prioritize candidate genes clinically related with the patient condition. Although this strategy allows POSTRE to more accurately predict both coding and long-range pathomechanisms, it also limits POSTRE applicability to certain diseases/phenotypes. However, this limitation should be reduced as functional genomic data becomes available for an increasing number of human cell types/tissues and is incorporated into POSTRE’s pipeline.

#### Multiple SVs as input

Considering the large number of SVs typically present in a human genome (around 10.000) (Lappalainen et al., 2019; Collins et al., 2020) as well as the increasing sizes of patient cohorts in which SVs are identified (Redin et al., 2017; Middelkamp et al., 2019), SV analysis tools should be ideally able to accept multiple SVs as input. However, not all the evaluated tools (Table 4) offer this possibility and can only analyze one SV at a time (e.g. TADeus2 and StrVCTVRE web browsers).

#### Interpretation of prediction results

Another main distinctive feature of POSTRE is that, in addition to predicting if a SV is pathogenic or not (e.g. Strvctvre (Sharo et al., 2022) or SVScore (Ganel et al., 2017)), it also provides a report explaining the genetic and molecular basis of such predictions in a particular cellular context. This information can significantly facilitate downstream experimental or *in silico* analyses to further characterize and validate the disease molecular mechanisms.

#### Required computational skills and User Experience

Tools designed to predict the pathogenicity of SVs or any other genetic alteration should be ideally usable by a broad scientific community, including scientists with limited computational skills. This is particularly relevant considering that the *in silico* interpretation of SVs might often involve the usage of not one, but several computational tools, as each tool has its own strengths and weaknesses (Richards et al., 2015). One way to improve tool usability is through the development of GUIs (graphical user interfaces). However, some of the current available SV prediction tools (e.g. SVScore, TADA) work through command line interfaces, thus complicating their usage. In contrast, once installed through a very simple procedure (i.e. running one command in a R environment), POSTRE can be executed with a user-friendly GUI that provides the user with detailed graphical and written reports for those SVs predicted as pathogenic.

## Discussion

SVs can affect gene function through either coding (e.g. gene deletions, gene fusions) or long-range (e.g. enhancer adoption, enhancer disconnection) mechanisms, which in turn can significantly contribute to human disease, phenotypic diversity and evolution (Feuk et al., 2006; Stankiewicz and Lupski, 2010; Spielmann et al., 2018; Real et al., 2020). SVs acting through coding mechanisms have been described for multiple human disorders (Nanni et al., 1999; Milunsky et al., 2008; Spielmann and Mundlos, 2016). In addition, recent advances in whole-genome sequencing are revealing that SVs with long-range pathological consequences might be also highly prevalent (Zhang and Lupski, 2015;Spielmann and Mundlos, 2016; Krude et al., 2021). However, the prediction of these long-range pathomechanisms is challenging, partly due to the still incomplete characterization of enhancers across different human cell types and our limited understanding of the factors controlling enhancer-gene communication. Consequently, there is a lack of tools for predicting the pathological effects of SVs considering both coding and long-range mechanisms. In this work, we have presented POSTRE, a user-friendly software that can be used to analyze SVs implicated in a broad range of congenital abnormalities.

In comparison with previous computational tools dedicated to the pathological analysis of SVs (see POSTRE benchmarking section for more details), POSTRE offers several advantages:

i. POSTRE is not restricted to CNVs, as it can handle all major types of SVs (i.e. deletions, inversions, duplications and translocations).
ii. POSTRE is capable of predicting both coding and long-range pathogenic effects and can distinguish between a wide variety of pathomechanisms (e.g. gene deletion, enhancer adoption, enhancer disconnection, enhancer deletion). Long-range mechanisms are predicted assuming that genes and enhancers located within the same TAD can effectively communicate with each other (Lupiáñez et al., 2016) and that developmental genes (i.e. broad polycomb domains in promoter regions) show high enhancer responsiveness (Kraft et al., 2019; Pachano et al., 2021). However, accumulating evidences indicate that additional factors (e.g. linear distance, CpG islands, promoter DNA methylation, type of core promoter elements) can also influence enhancer-gene compatibility within TADs (Arnold et al., 2017; Pachano et al., 2021; Ringel et al., 2021; Galouzis and Furlong, 2022; Zuin et al., 2022). As these factors are uncovered and characterized, they could be incorporated into POSTRE’s scoring system in order to further improve its specificity.
iii. POSTRE considers the cellular context to predict the pathological effects of SVs. To achieve this, POSTRE uses functional genomics information (e.g. gene expression levels, enhancer maps) generated in cell types/tissues deemed important for the patient phenotype (e.g. different brain developmental stages for neurodevelopmental defects). As a result, POSTRE not only predicts whether a SV is likely to be pathogenic but also the cellular context where such pathogenicity might be manifested, which can facilitate downstream experimental and/or *in silico* analyses. Furthermore, the independent evaluation of a given SV in various cell types/tissues offers the possibility of identifying either different candidate genes or different pathological mechanisms involving the same candidate gene depending on the cellular context (Spielmann et al., 2018). However, considering the cellular context can also introduce certain limitations, as illustrated by the patient described in Table1 in whom upregulation of *SHH* due to the duplication of its cognate ZRS enhancer causes limb abnormalities (Lohan et al., 2014). The ZRS enhancer drives *SHH* expression specifically within the ZPA (Hill and Lettice, 2013), which only represents a small fraction of all the limb cells during embryogenesis. Consequently, since POSTRE uses bulk genomic data generated in whole limb buds (Gerrard et al., 2016, 2020), the ZRS enhancer can not be identified and *SHH* appears as an inactive gene, thus precluding POSTRE from successfully predicting the effects of the patient duplication. In addition to the problem of using bulk data from heterogeneous tissues, the lack of genomic data for the appropriate developmental stages or differentiation time points may also affect the identification of pathologically relevant genes. Nevertheless, recent advances in single-cell genomics are expected to dramatically expand the catalogue of cell types and developmental stages in which gene expression profiles and enhancer maps can be explored (Abe et al., 2022; Eraslan et al., 2022; Tabula Sapiens Consortium* et al., 2022). Therefore, as single-cell datasets are generated in human embryos and incorporated to POSTRE, the tool will be capable of handling additional congenital abnormalities and also its sensitivity will increase.

One major discussion in the field of data sciences, particularly in the health care area, is model interpretability (Lipton, 2016; Reddy, 2022). Dealing with complex models complicates the user capacity to understand and learn from the predictions, which, in the biomedicine field, can limit the impact that such models can have on disease diagnosis or treatment (Vellido, 2020; Quinn et al., 2022). POSTRE’s scoring criterion is built on a set of simple and comprehensible rules based on the current knowledge about enhancers and the pathological relevance of genes, thus conceptually resembling the scoring criteria proposed in (Middelkamp et al., 2019) or TADeus2 (Poszewiecka et al., 2022). Hence, since the scoring criterion is simple, it is also easier to explain the results to the user. However, there is an increasing tendency towards using artificial intelligence to create prediction models. Currently, the standard option to create such models consists on providing a group of variables (predictors) measured in a set of observations and with a certain outcome associated to each observation (response variable). Next, during a training phase, a machine learning method automatically assigns a weight to each of the predictors and builds a model to predict the response variable (Nichols et al., 2019). During this phase, complex interactions among the predictors may be automatically established in the model, which complicates the capacity to interpret it afterwards. On top of that, if the training data does not faithfully recapitulate the general population, it can create biases in the algorithm criterion through overfitting (Nichols et al., 2019; Hertzberg et al., 2020), as it occurs with TADA (Hertzberg et al., 2020). Since databases used to train SV classifiers are biased towards coding deleterious effects, due to the scarcity of reported SV with long-range pathogenic effects, the classifiers can also suffer from the same bias and, thus, might not be appropriate for the prediction of non-coding (e.g. long-range) pathomechanisms (Hertzberg et al., 2020). In the case of POSTRE, since coding and long-range effects are evaluated independently (see Methods), the problem of coding effects dominating with respect to long-range effects is avoided, and overfitting in this context is minimized.

In summary, POSTRE is a user-friendly software to predict the pathomechanisms whereby SVs can cause congenital disorders. POSTRE can handle all major types of SVs, considers both coding and long-range mechanisms and performs its predictions in a cellular context-dependent manner. Altogether, these features make POSTRE rather unique among the SV prediction tools, particularly with respect to enhanceropathies, for which there is still a clear need of tools capable of linking non-coding variation with human disease (Claringbould and Zaugg, 2021).

## Acknowledgements

We would like to thank Maria Mariner Fauli for her advice on POSTRE’s graphical design and help with the elaboration of some figures. We would also like to thank Magdalena Laugsch, Julia Baptista, Ayat Essabi and all the Rada-Iglesias lab members for insightful comments and suggestions about POSTRE and POSTRE’s manuscript.

## Author Contributions

V.S.G. developed POSTRE and performed all the analysis. V.S.G and A.R.I conceptualized the project. A.R.I supervised the work. V.S.G and A.R.I wrote, the manuscript. All the authors read and approved the final manuscript.

## Materials and Methods

### POSTRE software installation

POSTRE is built with the Shiny framework (Chang et al., 2021) and, thus, the majority of its code is written in R. Regarding the user interface, it has been developed using Shiny libraries and custom html, css and javascript code.

The full set of instructions (including videos and tutorials) explaining how to download and run POSTRE is provided in GitHub https://github.com/vicsanga/Postre. Briefly, the user needs to install R (version >=3.5.0) and then simply run the following command in the R console: source(“https://raw.githubusercontent.com/vicsanga/Postre/main/Postre_wrapper.R”)

### POSTRE main functionalities

- **Single SV Submission:** Allows the user to submit one SV and analyze it in the context of one phenotype. If pathogenic events are found, POSTRE provides an explanatory report. This report contains a set of graphics and text that facilitate the interpretation of the SV pathogenicity. It also provides links to different external resources (e.g. OMIM (Amberger et al., 2009), Medgen (Louden, 2020) and the Mouse Genome Database (Bult et al., 2019) containing information about the gene/s affected by the SV. Lastly, the user can also visualize disease-relevant genomic data within the affected locus through the UCSC genome browser (Kent et al., 2002).
- **Multiple SVs Submission:** Allows the user to submit multiple SVs simultaneously and to assign multiple phenotypes to each of them. All the downstream predictions are summarized in a set of tables that classify the detected pathogenic events by phenotype, gene, and type of pathological mechanism (i.e. coding LOF, coding GOF, long-range LOF and long-range GOF). Next, a detailed report for each of the submitted SVs can be easily obtained by executing a Single Submission job for the SV of interest.
- **Explore Previous SVs:** Allows the user to navigate POSTRE predictions for patient SVs reported in public databases. The predicted pathogenic events are presented through different tables that include information about phenotypes, genes, and type of pathological mechanism (i.e. coding LOF, coding GOF, long-range LOF and long-range GOF). The detailed report for each of the analyzed SVs can be easily obtained by executing a Single Submission job for the SV of interest.

### TAD maps annotation

POSTRE uses TADs as a proxy of gene regulatory domains in order to predict the long-range pathomechanisms of SVs. All genes found within the TAD/s affected by a SV are initially considered as potentially involved in disease pathogenesis (candidate genes). Depending on the patient phenotype, POSTRE uses TAD maps generated in various cell types/tissues (Supplementary Data 1). These TAD maps were obtained from publically available repositories through different approaches (Supplementary Data 1):

- TAD coordinates already available (no processing needed).
- TAD coordinates inferred from available TAD boundary maps. In this case, TADs are defined as the regions located between neighboring TAD boundaries.
- TAD coordinates obtained with the DomainCaller tool (https://github.com/XiaoTaoWang/domaincaller/) and the 50kb contact matrices provided in .hiC files.

### LOF and GOF definitions

The terms LOF and GOF appear recurrently along the manuscript. To avoid misunderstandings, it is important to clarify that these terms are used as follows:

- LOF (Loss of Function): pathogenic loss of gene function or expression through either coding or non-coding mechanisms.
- GOF (Gain of Function): pathogenic gain of gene function or expression through either coding or non-coding mechanisms.

### POSTRE pathogenicity score

Firstly, the breakpoints of the SVs are mapped in the context of the TAD maps associated with the phenotype of interest. Once the affected TADs (i.e. regulatory domains) are determined, all the genes located within them are selected as potential candidates associated with the patient disease. Subsequently, POSTRE integrates a set of genetic and genomic features in order to assign a pathogenicity score (PS) to each candidate gene considering in parallel both LOF and GOF scenarios. These gene PS are independently calculated in each of the cell types/tissues considered for a given phenotype. In addition, POSTRE offers the possibility to compute the PS using two alternative running modes (i.e. Standard and High-Specificity; see details below). Finally, all candidate genes are ranked according to their PS in each of the considered cell types/tissues.

The features used to calculate the gene PS can be broadly divided into the following categories or sub-scores:

- Gene-phenotype relationships (*genePhenoScore*)
- Gene features (*geneFeaturesScore*)

- Dosage sensitivity (*dosageSensitivityScore*)
- Breadth of polycomb domains in promoter regions (*polycombScore*)
- Gene expression (*geneExpressionScore*)
- Enhancer activity (*geneEnhancerScore*)

#### Gene-Phenotype relationships: Computing the genePhenoScore

The *genePhenoScore* metric is used to quantify the relationship between the candidate genes and the SV associated phenotype. Before describing how the *genePhenoScore* was calculated, we briefly explain how the associations between genes and phenotypes were established.

Firstly, we obtained a collection of Human Phenotype Ontology (HPO) terms (Köhler et al., 2021) for each of the considered phenotypic categories (*i.e.* Cardiovascular, Head-Neck, Limbs and Neurodevelopmental). HPO terms represent standardized phenotypic categories that facilitate the annotation of clinical abnormalities. HPO terms are organized following a hierarchical and nested structure. For example, the HPO term Absent speech (HP:0001344) belongs, following a hierarchical order, to the *Delayed speech and language* (HP:0000750), *Neurodevelopmental abnormality* (HP:0012759), *Abnormality of the nervous system* (HP:0000707) and *Phenotypic abnormality* (HP:0000118) categories. In POSTRE, a set of reference HPOs, together with all their nested and more specific terms, were selected for each of the considered phenotypes. Following the previous example, when considering neurodevelopmental phenotypes we used *Neurodevelopmental abnormality* (HP:0012759) as a reference HPO, which, among others includes the *Delayed speech language* (HP:0000750) and *Absent speech* (HP:0001344) HPO terms. In this regard, the reference HPOs selected for the phenotypes considered in POSTRE were:

- Cardiovascular: *Abnormality of the cardiovascular system* (HP:0001626)
- Head-Neck: *Abnormality of head or neck* (HP:0000152)
- Limbs: *Abnormality of limbs* (HP:0040064)
- Neurodevelopmental: *Neurodevelopmental abnormality* (HP:0012759)

To retrieve all the HPO terms associated with each of the reference HPOs, we used the *hp.obo* file from the Downloads/Ontology section in the HPO website (Köhler et al., 2021).

On the other hand, human genes were also annotated according to HPO terms. For each gene, all its associated HPO terms were retrieved using the *genes_to_phenotype.txt* file from the Downloads/Annotation section in the HPO website (Köhler et al., 2021). Next, genes associated with at least one of the HPO terms present within the reference HPO phenotypic groups described above were linked with the main phenotypes considered by POSTRE (i.e. Cardiovascular, Head-Neck, Limbs or Neurodevelopmental). For example, if a gene is only associated with the *Delayed speech language* (HP:0000750) term, it will still be linked with the Neurodevelopmental phenotype because it is contained within the more general *Neurodevelopmental abnormality* (HP:0012759) HPO category.

A similar approach was used to associate human genes with mouse phenotypes as defined in the Mammalian Phenotype Ontology (MPO) (Smith et al., 2005). MPO terms, which are equivalent to HPOs, also present a hierarchical and nested organization. As described for HPOs, a set of reference MPOs and all their nested and more specific terms were selected for each of the considered phenotypes:

- Cardiovascular: *Cardiovascular system phenotype* (MP:0005385)
- Head Neck: *Craniofacial phenotype* (MP:0005382) and *abnormal neck morphology* (MP:0012719)
- Limbs: *Limbs/digits/tail phenotype* (MP:0005371)
- Neurodevelopmental: *Behavior/neurological phenotype* (MP:0005386)

To retrieve all the MPO terms associated with each of the reference MPOs, we used the mp.obo file from the OBO Foundry website (Jackson et al., 2021). Next, to link human genes with MPO terms, we used the *HMD_HumanPhenotype.rpt* file from the Mouse Genome Database (Bult et al., 2019). Lastly, human genes associated with at least one of the MPO terms present within the reference MPO phenotypic groups described above were linked with the main phenotypes considered by POSTRE (i.e. Cardiovascular, Head-Neck, Limbs or Neurodevelopmental).

The previous relationships between human genes and human/mouse phenotypes were used to compute the *genePhenoscore.* When considering the phenotype associated with a SV (POSTRE input), a candidate gene receives a *genePhenoscore*=1 if it is associated with the same phenotype in humans, a *genePhenoscore*=0.5 if it is associated with the same phenotype in mice but not in humans (consideration only applied in the *Standard* Running Mode) and a *genePhenoScore*=0 otherwise.

#### Dosage sensitivity: computing the dosageSensitivityScore

The *dosageSensitivityScore* is used to quantify the dosage sensitivity of the candidate genes. Haploinsufficient (HI) genes tend to be associated with disease not only when present as a single copy (or downregulated), but also when present in supernumerary copies (or upregulated) (Collins et al., 2020). HI gene scores were retrieved from two different sources: (i) HI scores ranging between 0 (low HI) and 1 (High HI) were obtained from (Huang et al., 2010; Lek et al., 2016); (ii) dosage sensitivity information was obtained from ClinGen (ftp://ftp.clinicalgenome.org/ClinGen_haploinsufficiency_gene_GRCh37.bed) (Rehm et al., 2015) and genes with strong evidence for dosage sensitivity (ClinGen score = 3) were assigned a HI score=1. Then, between these two alternative HI scores, the highest one was selected for each gene. Next, genes with HI scores >= 0.75 were assigned such value as their *dosageSensitivityScore*, while genes with HI scores < 0.75 were assigned a *dosageSensitivityScore* = 0.

#### Breadth of polycomb domains in promoter regions: computing the polycombScore

The *polycombScore* was used to quantify the breadth of polycomb domains in gene promoters. Previous work showed that genes whose promoters are covered by broad polycomb/H3K27me3 domains when inactive largely correspond with major developmental genes frequently implicated in congenital disorders (Rehimi et al., 2016; Shim et al., 2020). Moreover, these developmental genes display high enhancer responsiveness (Kraft et al., 2019; Pachano et al., 2021). The *polycombScore* ranges from 0 (no or narrow polycomb domain) to 1 (broad polycomb domain).

To compute the *polycombScore* two different approaches were considered.

- Approach#1: H3K27me3 ChIP-seq data generated in: ESCs (GSE24447; H3K27me3: SRR067372, input:SRR067371), Neural Crest Cells (GSE108518; H3K27me3: SRR6418921, input: SRR6418919) and Cardiomyocytes (GSE85628; H3K27me3: SRR4032228, input: SRR4032231) were used independently to call H3K27me3 peaks using MACS2 (Feng et al., 2012) with the broad peak calling mode. Peaks with a fold-enrichment > 3 and Q-value < 0.1 were considered. Subsequently, peaks within 1 kb of each other were merged using *bedtools* (Quinlan and Hall, 2010), and associated with a protein-coding gene when overlapping a TSS. Lastly, the knee of the size distribution of the H3K27me3 peaks associated with genes was determined with *findiplist()*(inflection R package; https://cran.r-project.org/web/packages/inflection/vignettes/inflection.html). Upon curvature analysis, genes with H3K27me3 peaks > 6.6 kb in any of the considered cell types were assigned a *polycombScore*=1 (i.e. broad polycomb domains) and all other genes received a *polycombScore*=0.
- Approach#2: It is based on the repressive tendency score (RTS) described in (Shim et al., 2020), a metric that quantifies the association between genes and broad H3K27me3 domains. This metric is computed considering data generated in hundreds of different cell types. RTS ranges from 0 (no or narrow polycomb domain) to 1 (broad polycomb domain). Based on RTS, Shim et al., 2020 defined three groups of genes: (i) “regulatory” genes (high RTS), (ii) “structural” genes (medium RTS) and (iii) “housekeeping” genes (low RTS). “Regulatory” genes were assigned a *polycombScore*=1 and “housekeeping genes” were assigned a *polycombScore*=0. For the “structural” genes we applied a 0-1 min-max normalization, considering as the minimum value the minimum RTS of all genes and as the maximum value the smallest RTS among the regulatory genes. Thus, for the structural genes:

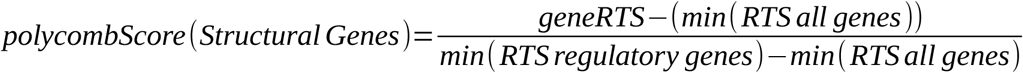

Finally, the highest *polycombScore* obtained among these two different approaches was assigned to each gene.

#### Gene expression: computing the geneExpressionScore

The *geneExpressionScore* is used to incorporate gene expression levels into POSTRE’s PS. A detailed list of the gene expression datasets considered for the different phenotypes, together with the methodologies used to quantify gene expression in each dataset, is provided in Supplementary Data 1.

Considering the expression status of the candidate genes is particularly relevant for loss of function (LOF) situations, as the disease-causative genes must be expressed in the relevant cell types/tissues. We defined a minimum (FPKM=1) and a maximum expression threshold (FPKM=10): genes with FPKM<1 are considered as not expressed and are assigned a *geneExpressionScore*=0; genes with FPKM>10 are considered to be expressed and are assigned a *geneExpressionScore*=1; genes with 1<FPKM<10 are subject to a 0-1 min-max normalization to determine its *geneExpressionScore*, considering as min-max values the expression thresholds.

#### Computing the geneFeatureScore

This metric results from the aggregation of the *geneExpressionScore*, *polycombScore* and the *dosageSensitivityScore*.

For this metric to be > 0 either the *polycombScore* or the *dosageSensitiveScore* must be >0.75 in the *Standard* running mode, while both scores must be >0.75 in the *High-Specificity* running mode. The *geneFeatureScore* is computed as follows:

<u>When evaluating LOF:</u>

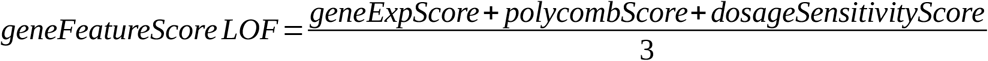

<u>When evaluating GOF:</u>

GOF pathomechanisms can occur through (i) upregulation of already active genes through duplications directly affecting the genes or some of their cognate enhancers; (ii) upregulation of either active or inactive genes through interactions with ectopic enhancers (i.e. enhancer adoption). The *geneFeatureScore* is computed differently for these two GOF scenarios:

- Duplications directly affecting active genes or some of their cognate enhancers:

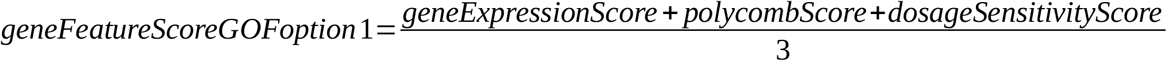
- SVs leading to enhancer adoption by either active or inactive genes:

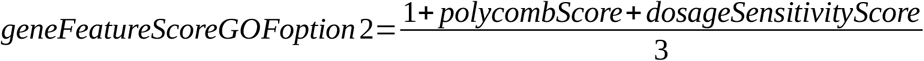

For GOF involving enhancer adoption we deemed as irrelevant whether the candidate genes are initially active or inactive and, thus, the *geneExpressionScore*=1.

#### Changes in enhancer activity: computing the geneEnhancerScore

The *geneEnhancerScore* quantifies changes in enhancer activity between Control and Patient alleles. Firstly, enhancers are annotated in the different cell types/tissues considered by POSTRE. To annotate enhancers, we considered H3K27ac peaks located at least 10kb away from any protein-coding gene TSS. Depending on the cell type/tissue, H3K27ac peaks were either already available or were called using MACS2 (Feng et al., 2012). For some cell types/tissues, chromatin accessibility (e.g. DNAse-seq, ATAC-seq) data and/or ChIP-seq data for other regulatory proteins (e.g. p300) were available in addition to H3K27ac. In those cases, enhancers were identified as regions showing overlapping peaks for H3K27ac and other available proteins/chromatin accessibility. The ChIP-Seq datasets used to annotate enhancers in the different cell types/tissues together with the methodologies applied in each case are more extensively described in Supplementary Data 1. In addition, H3K27ac ChIP-seq bigwig files were also obtained for each of the relevant cell types/tissues. These H3K27ac bigwig files were either already available in public repositories or generated with *deepTools* (Ramírez et al., 2016) using *bamCoverage* (reads per genomic content, RPGC, normalization). Next, for each cell type/tissue, the enhancer lists and H3K27ac bigwig files were combined to quantify enhancer activity as the maximum H3K27ac levels for each enhancer using the *bigWigAverageOverBed* UCSC binary tool. As a result, each enhancer was assigned an *enhancerIndividualActivity* score according to the following formula:

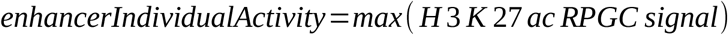

Then, for each candidate gene an *enhancerActivityControl* metric is calculated as the sum of the *enhancerIndividualActivity* scores assigned to the enhancers located within the Control (i.e. without any SV) candidate gene regulatory domain. Likewise, an *enhancerActivityPatient* metric is calculated as the sum of *enhancerIndividualActivity* scores assigned to the enhancers located within the rearranged candidate gene regulatory domain. We assume that individual enhancer activities are not affected by the SVs. Next, for each candidate gene the ratio between the *enhancerActivityPatient* and *enhancerActivityControl* values are computed assuming either LOF or GOF scenarios:

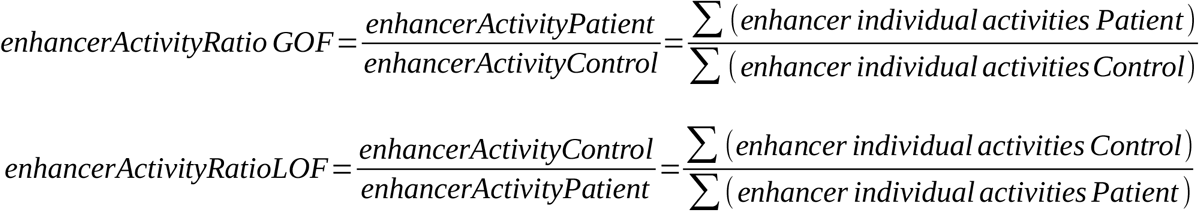

For LOF (*enhancerActivityRatioLOF)*, the *enhancerActivityPatient* value is computed only considering the cognate enhancers that remain within the rearranged candidate gene regulatory domain. For GOF (*enhancerActivityRatioGOF),* the *enhancerActivityPatient* value is computed considering both the remaining cognate enhancers as well as the ectopic enhancers located within the rearranged candidate gene regulatory domain.

*enhancerActivityRatioGOF* >1 indicates that the candidate gene shows higher enhancer activity in the Patient regulatory domain. *enhancerActivityRatioLOF* >1 indicates that the candidate gene shows higher enhancer activity in the Control/wild-type regulatory domain.

Lastly, *enhancerActivityRatios* are transformed into 0-1 values to obtain the final *genEnhancerScore* used by POSTRE to compute the PS, with higher *genEnhancerScores* indicating more relevant changes in enhancer activity. In order to make these 0-1 transformations, we used different thresholds for the *enhancerActivityRatioLOF* (*maxLOFratio=1.2)* and *enhancerActivityRatioGOF (maxGOFratio=2)* ratios. These thresholds were estimated based on the analyses of SVs with experimentally validated pathomechanisms (i.e. positive controls). Therefore, the final *geneEnhancerScores* for each candidate gene were computed as follows:

- *geneEnhancerScoreLOF*: If *enhancerActivityRatioLOF* ≥ *maxLOFratio*, then *geneEnhancerScoreLOF*=1. If *enhancerActivityRatioLOF* is ≤ 1 then *geneEnhancerScoreLOF*=0. If 1 < *enhancerActivityRatioLOF* < *maxLOFratio*, then a min-max 0-1 normalization is applied.
- *geneEnhancerScoreGOF*: If *enhancerActivityRatioGOF*≥ *maxGOFratio*, then *geneEnhancerScoreGOF*=1. If *enhancerActivityRatioGOF* is ≤ 1 then *geneEnhancerScoreGOF*=0. If 1 < *enhancerActivityRatioGOF* < *maxGOFratio*, the a min-max 0-1 normalization is applied.

#### Calculating the pathogenic score (PS)

To calculate the PS, the metrics described in previous sections are integrated. For each candidate gene and cell type/tissue analyzed, PS are computed assuming both LOF and GOF scenarios. Moreover, PS are computed differently depending on whether or not the SVs directly affect gene sequences.

#### I. Long-range pathomechanisms: candidate genes not directly affected by the SV

<u>PS for LOF</u>

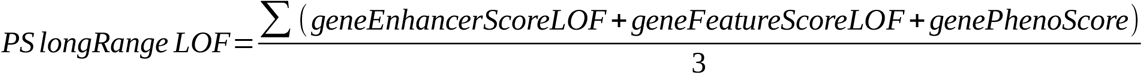

- NOTE: candidate genes whose expression < 1 FPKM (minimum expression threshold) are considered as not expressed and their PS score=0.

<u>PS for GOF</u>

If the SV is predicted to cause ectopic interactions between the candidate gene and non-cognate enhancers (e.g. through TAD fusion or TAD shuffling), the *geneExpressionScore* is not considered relevant:

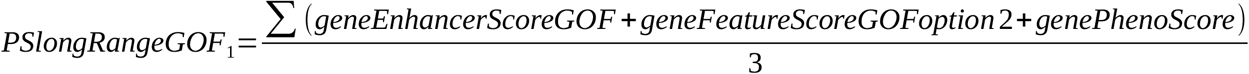

If the SV is not predicted to cause ectopic interactions between the candidate gene and non-cognate enhancers (e.g. intra-TAD duplication), the *geneExpressionScore* is considered important:

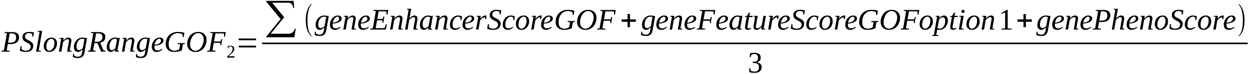

#### II. Coding pathomechanisms: candidate genes directly affected (deleted, truncated or duplicated) by the SV

If the candidate gene is deleted or truncated, only LOF will be evaluated and the PS for GOF will be set to 0. If the gene is duplicated, only GOF will be assessed and the PS for LOF will be set to 0.

<u>PS for LOF</u>

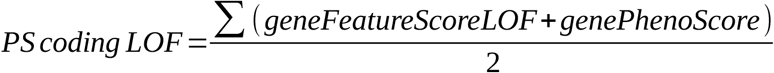

- NOTE: candidate genes whose expression < 1 FPKM (minimum expression threshold) are considered as not expressed and their PS score=0.

<u>PS for GOF</u>

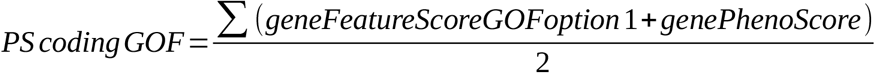

- NOTE: candidate genes whose expression < 1 FPKM (minimum expression threshold) are considered as not expressed and their PS score=0.

If the SVs cause a gene deletion or truncation, then long-range pathomechanisms are not considered for those candidate genes. However, if the SVs cause gene duplication/s, long-range pathomechanisms involving enhancer adoption and the formation of new TADs can still occur (Franke et al., 2016; Spielmann et al., 2018). Hence, for SVs resulting in gene duplications, coding and long-range GOF PS are computed and the highest one is selected as the most likely pathomechanism.

Given all the previous PS models, POSTRE applies them as follows:

1. POSTRE evaluates whether the SV directly affects the candidate gene sequence “coding” or not “long-range” alteration.
2. Based on the impact of the SV over the candidate gene, the corresponding GOF and LOF PS models are applied.
3. Each candidate gene gets two PS scores: *PS_GOF* and *PS_LOF*. If any of these two PS scores is higher than a pathogenic threshold (0.8), then a detailed report describing the predicted pathomechanism is provided.

## Additional tools and resources

The genomic data displayed in the UCSC genomic browser that is accessible through POSTRE reports is hosted either at the CyVerse Discovery Environment (https://de.cyverse.org), or at the GEO database (Barrett et al., 2013).

## Reference genome

The reference genome currently handled by POSTRE is hg19. Accordingly, all genomic coordinates provided in the supplementary material correspond with this reference genome.

## Supplementary Figures

**Supplementary Figure 1.**
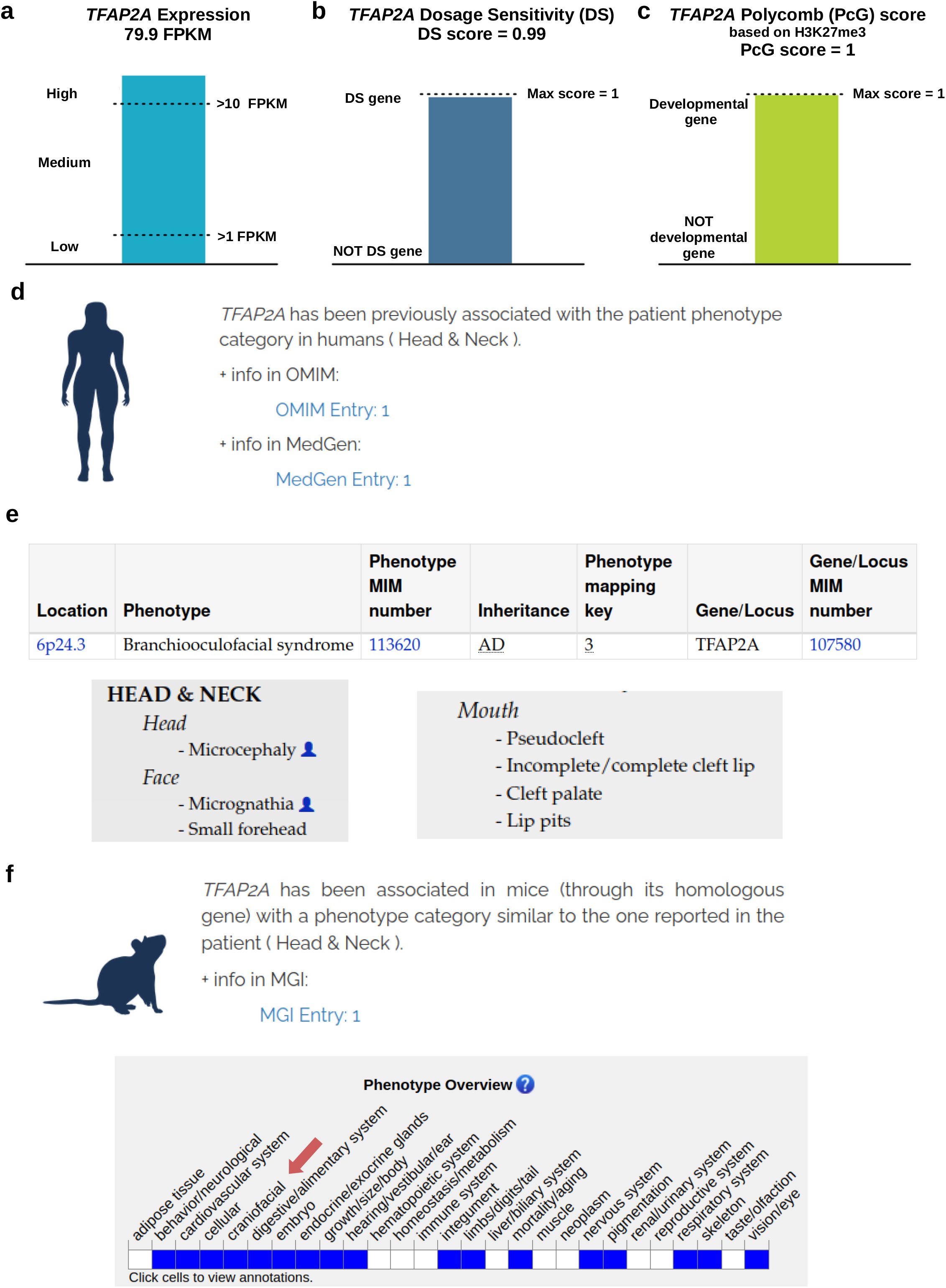
Additional information provided by POSTRE for the *TFAP2A* patient. **a.** *TFAP2A* expression levels measured in NCC (NeuralCrestLate) by RNA-seq are shown as FPKMs. **b**, *TFAP2A* displays a high Dosage Sensitivity (DS) Score, suggesting that humans are likely to be haploinsufficient for this gene. **c**, *TFAP2A* received a high Polycomb (PcG) Score since, when inactive, it is embedded within a broad PcG domain. **d**, *TFAP2A* is associated with craniofacial (Head&Neck) abnormalities in humans according to information found in OMIM and MedGen databases. **e**, Summary of the craniofacial phenotypes associated with *TFAP2A* according to OMIM (OMIM Entry 1). **f**, *TFAP2A* homologous gene in mice (*Tfap2a)* is associated with craniofacial abnormalities according to information found in the MGI database.

**Supplementary Figure 2.**
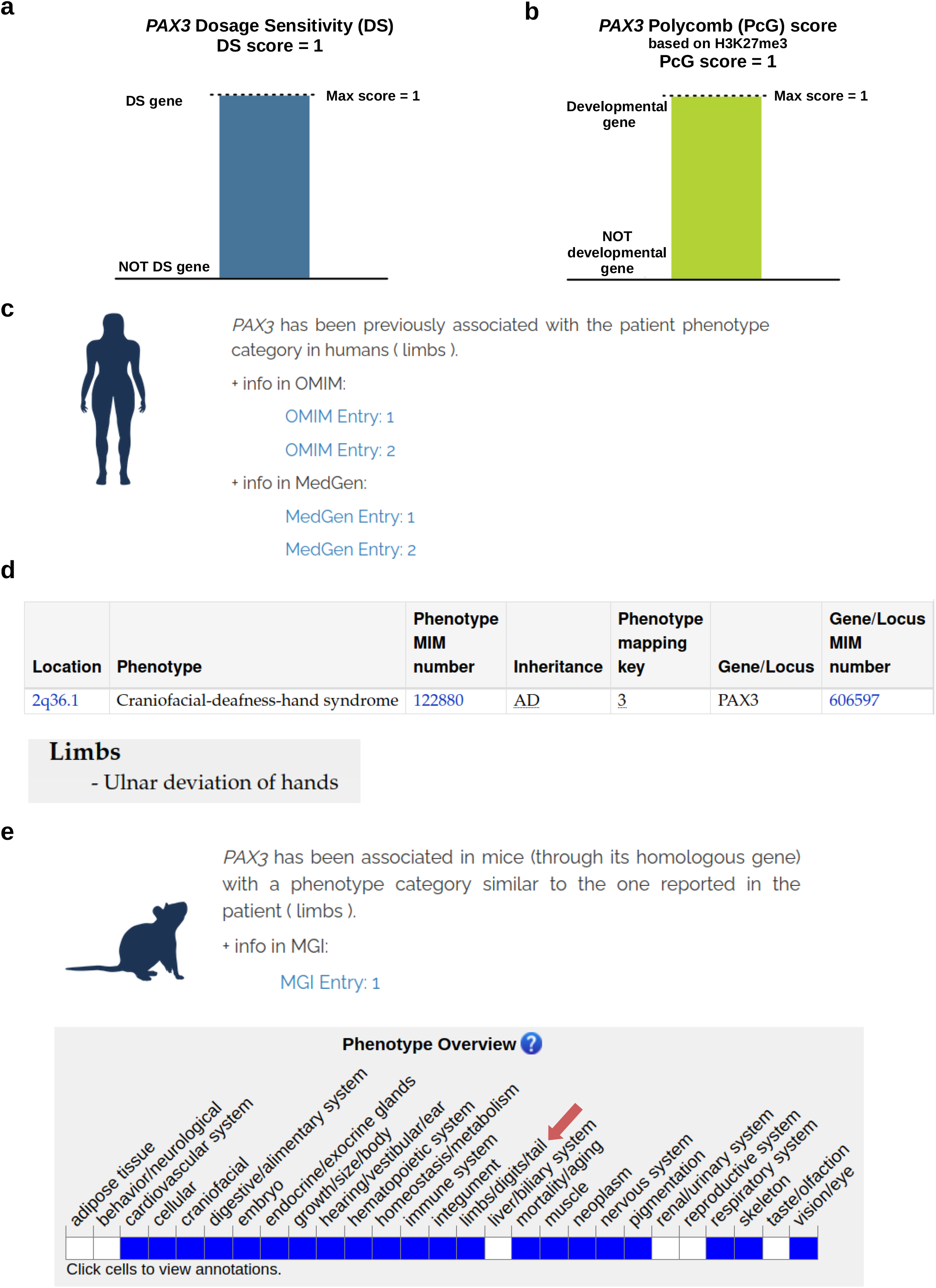
Additional information provided by POSTRE for the *PAX3* patient. **a.** *PAX3* displays a high Dosage Sensitivity (DS) Score, suggesting that humans are likely to be sensitive to changes in its copy number or expression levels. ***b*,** *PAX3* received a high Polycomb (PcG) Score since, when inactive, it is embedded within a broad PcG domain. **c**, *PAX3* is associated with limb abnormalities in humans according to information found in OMIM and MedGen databases. **d**, Summary of the limb phenotypes associated with *PAX3* according to OMIM (OMIM Entry 1). **e**, PAX3 homologous gene in mice (*Pax3)* is associated with limb abnormalities according to information found in the MGI database.

**Supplementary Figure 3.**
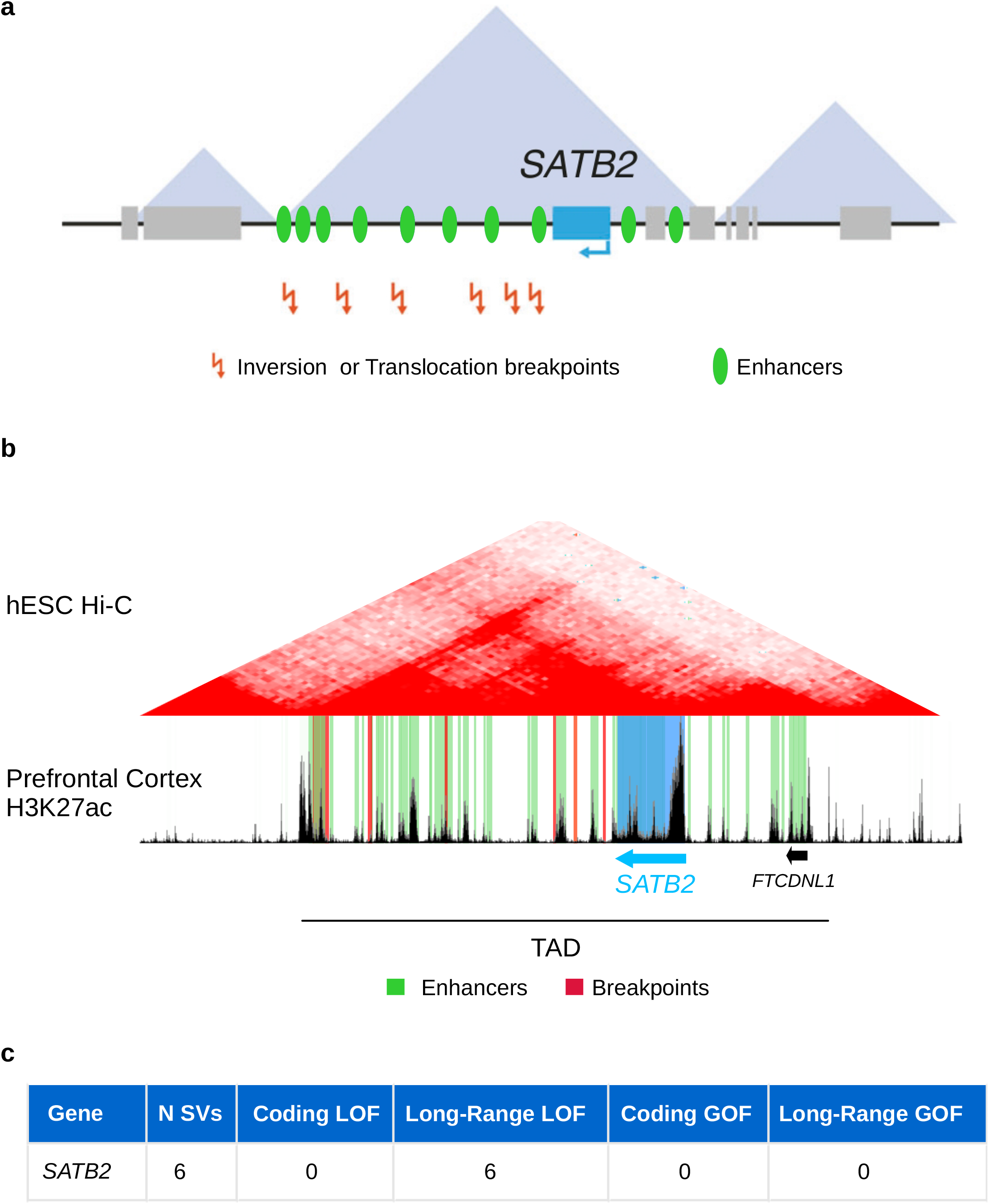
POSTRE predictions for SVs identified in patients with neurodevelopmental defects and predicted to affect SATB2 expression. **a**, Graphical representation of the *SATB2* locus (adapted from (D’haene and Vergult, 2021)) showing the genomic location of (i) six different SV breakpoints (inversions or translocations) found in patients with neurodevelopmental deffects (Supplementary Data 4) and (ii) active enhancers identified in embryonic brain prefrontal cortex. **b**, Genome browser view of the *SATB2* TAD. The breakpoints of the six different SVs are highlighted in red and active enhancers identified in the embryonic brain prefrontal cortex gestation week 18 (PfcGw18) based on H3K27ac ChIP-seq data are highlighted in green. **c**, Summary of the results obtained with POSTRE upon analysis of the six SVs described in (a). The six SVs are predicted to cause *SATB2 LOF* through a long-range (i.e. enhancer disconnection) pathomechanism.

**Supplementary Figure 4.**
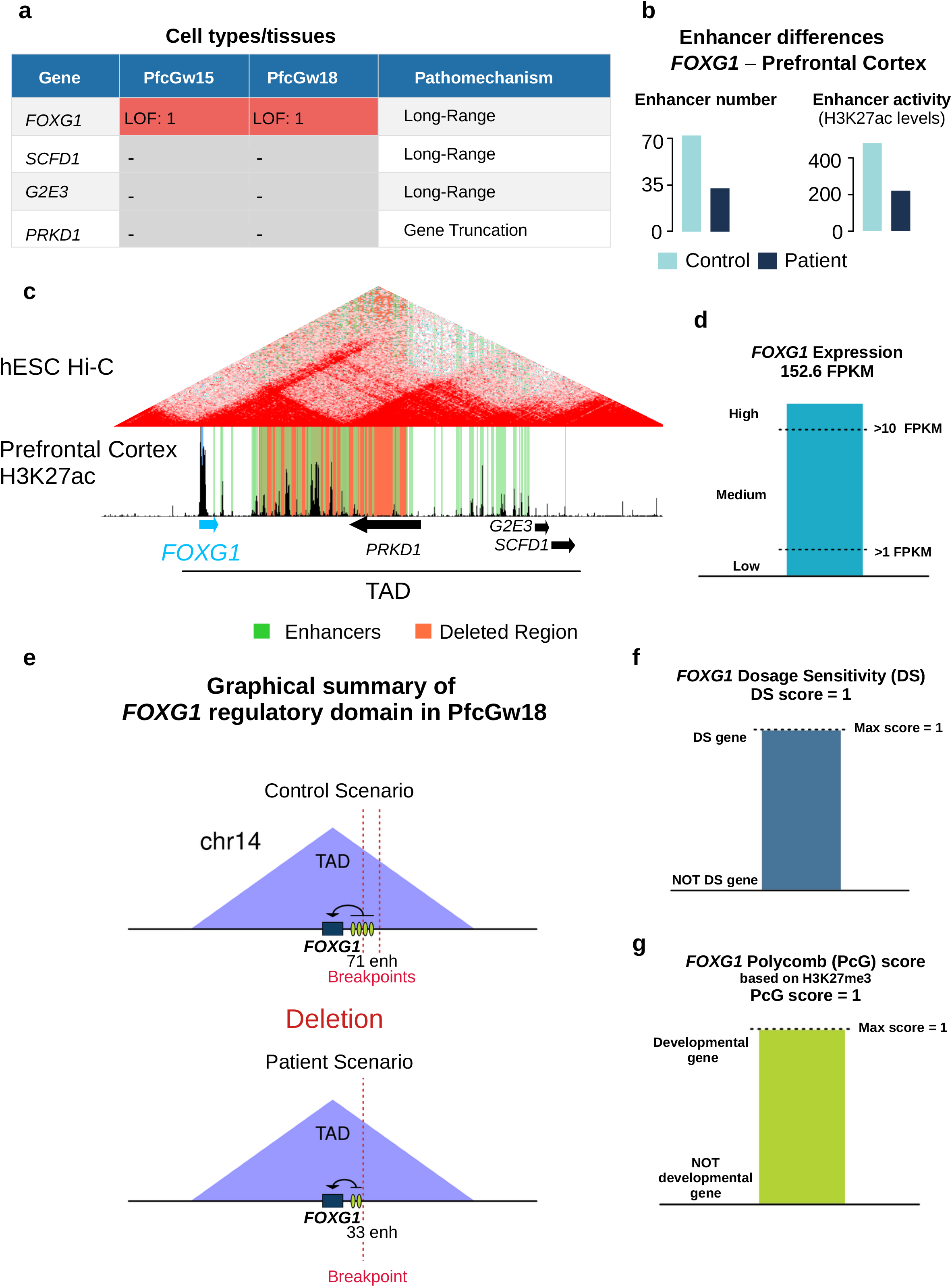
POSTRE results for one of the *FOXG1* patients. **a,** A deletion in Chr14 was previously reported for a patient with neurodevelopmental defects (*FOXG1* patient Nr 8 in Supplementary Data 4). POSTRE identified *FOXG1* as the causative gene in this patient and predicted that the deletion could lead to a loss of *FOXG1* expression through a long-range mechanism (enhancer deletion) in the brain prefrontal cortex gestation week 15 (PfcGw15) and 18 (PfcGw18). **b**, Enhancer activity (H3K27ac levels) and number of enhancers associated with *FOXG1* in embryonic brain prefrontal cortex (PfcGw18) are shown in the absence (Control) or presence (Patient) of the deletion. **c**, Genome browser view of the TAD affected by the deletion. The deletion is highlighted in orange and the prefrontal cortex active enhancers in PfcGw18 in green. **d.** *FOXG1* expression levels measured in the prefrontal cortex (PfcGw18) by RNA-seq are shown as FPKMs. **e**, *FOXG1* displays a high Dosage Sensitivity (DS) Score, suggesting that humans are likely to be haploinsufficient for this gene. *f*, *FOXG1* received a high Polycomb (PcG) Score since, when inactive, it is embedded within a broad PcG domain. **g**, Graphical abstract illustrating the changes in the regulatory landscape of *FOXG1* in PfcGw18 due to the deletion. The green ovals represent enhancers (enh).

**Supplementary Figure 5.**
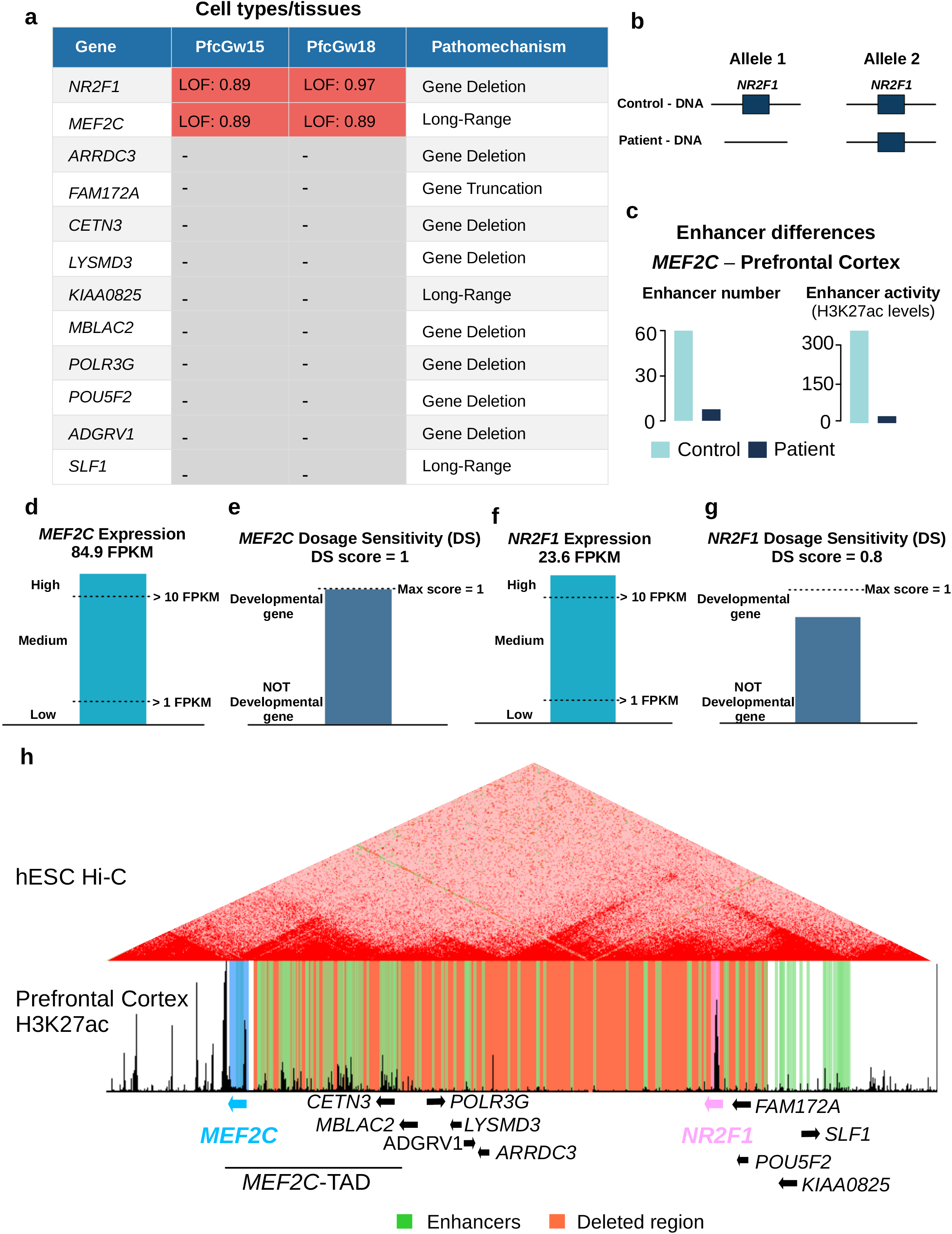
POSTRE results for one of the *MEF2C* patients. **a,** A deletion in Chr5 was previously reported for a patient with neurodevelopmental defects (*MEF2C* patient Nr 11 in Supplementary Data 4). POSTRE identified *NR2F1* and *MEF2C* as the two causative genes in this patient in the brain prefrontal cortex gestation week 15 (PfcGw15) and 18 (PfcGw18). For *MEF2C,* POSTRE predicted that the deletion could lead to a loss of *MEF2C* expression through a long-range mechanism (enhancer deletion). **b,** In addition, POSTRE also predicted *NR2F1* as a causative gene through a coding pathomechanism, since the deletion eliminates one of the two *NR2F1* alleles. **c**, Enhancer activity (H3K27ac levels) and number of enhancers associated with *MEF2C* in embryonic brain prefrontal cortex (PfcGw18) are shown in the absence (Control) or presence (Patient) of the deletion. **d and f,** *MEF2C* (d) and *NR2F1* (f) expression levels measured in the prefrontal cortex (PfcGw18) by RNA-seq are shown as FPKMs. **e and g**, *MEF2C* (e) and *NR2F1* (g) display high Dosage Sensitivity (DS) Scores, suggesting that humans are likely to be haploinsufficient for these genes. **h**, Genome browser view of the genomic region affected by the deletion, including the *MEF2C TAD*. The deletion is highlighted in orange and the prefrontal cortex active enhancers in PfcGw18 in green.

**Supplementary Figure 6.**
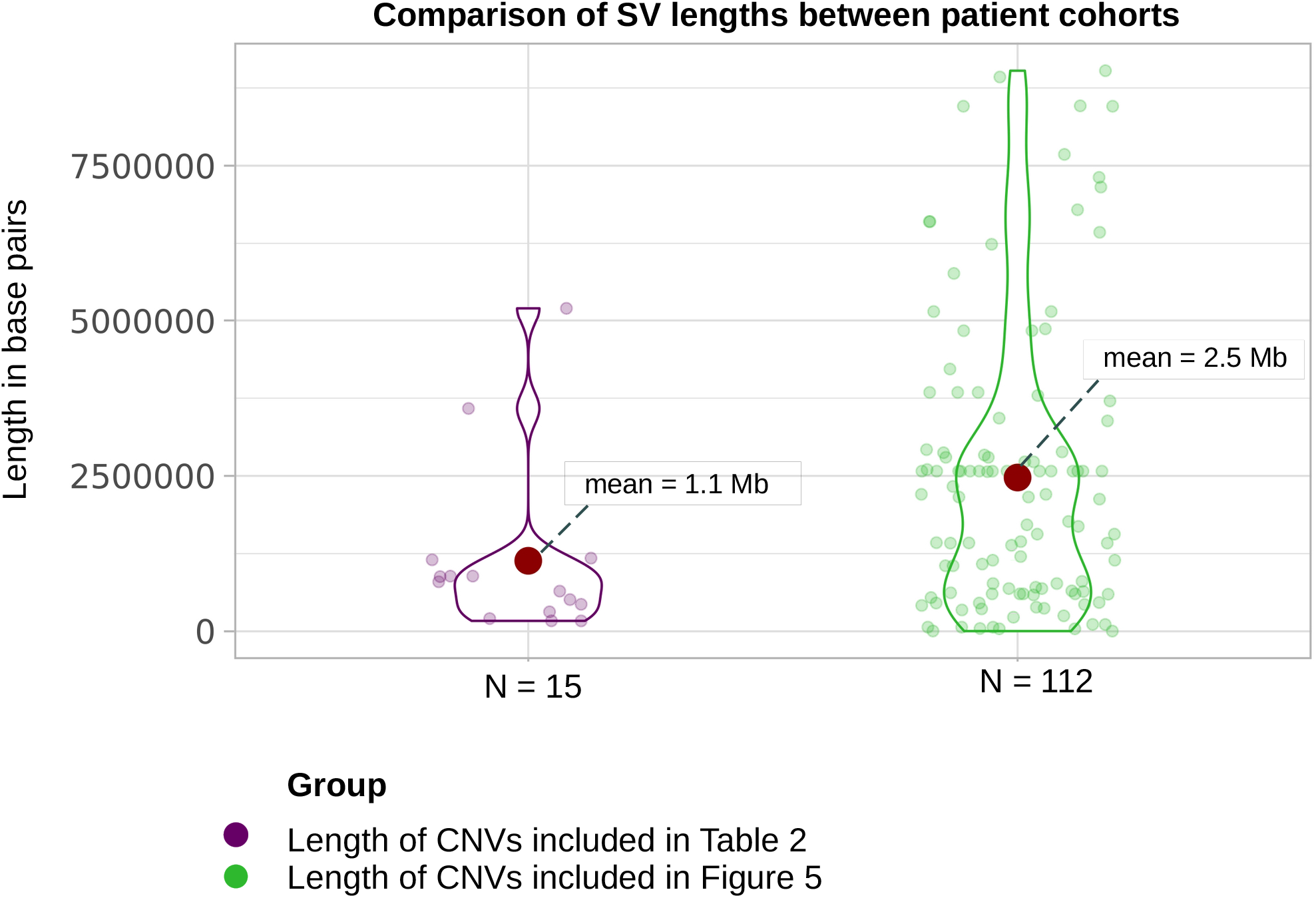
Comparison of SVs lengths between different patient cohorts. Violin plots showing the lengths of the SVs from the patient cohorts analyzed in Table 2 (purple, left) and Figure 5 (green, right). Each dot corresponds to a SV. Only CNVs (i.e. deletions and duplications) are considered.

## References

Abe, Y., Sakata-Yanagimoto, M., Fujisawa, M., Miyoshi, H., Suehara, Y., Hattori, K., et al. (2022). A single-cell atlas of non-haematopoietic cells in human lymph nodes and lymphoma reveals a landscape of stromal remodelling. Nat. Cell Biol. 2022 244 24, 565–578. doi:10.1038/s41556-022-00866-3.

Amberger, J., Bocchini, C. A., Scott, A. F., and Hamosh, A. (2009). McKusick’s Online Mendelian Inheritance in Man (OMIM®). Nucleic Acids Res. 37, D793. doi:10.1093/NAR/GKN665.

Arnold, C. D., Zabidi, M. A., Pagani, M., Rath, M., Schernhuber, K., Kazmar, T., et al. (2017). Genome-wide assessment of sequence-intrinsic enhancer responsiveness at single-base-pair resolution. Nat. Biotechnol. 35, 136–144. doi:10.1038/nbt.3739.

Barik, S., Pandita, N., Paul, S., Kumari, O., and Singh, V. (2021). Prevalence of congenital limb defects in Uttarakhand state in India – A hospital-based retrospective cross-sectional study. Clin. Epidemiol. Glob. Heal. 9, 99–103. doi:10.1016/j.cegh.2020.07.007.

Barnett, C. P., Mencel, J. J., Gecz, J., Waters, W., Kirwin, S. M., Vinette, K. M. B., et al. (2012). Choreoathetosis, congenital hypothyroidism and neonatal respiratory distress syndrome with intact *NKX2-1*. Am. J. Med. Genet. Part A 158A, 3168–3173. doi:10.1002/ajmg.a.35456.

Barrett, T., Wilhite, S. E., Ledoux, P., Evangelista, C., Kim, I. F., Tomashevsky, M., et al. (2013). NCBI GEO: Archive for functional genomics data sets - Update. Nucleic Acids Res. 41, D991– D995. doi:10.1093/nar/gks1193.

Batut, P. J., Bing, X. Y., Sisco, Z., Raimundo, J., Levo, M., and Levine, M. S. (2022). Genome organization controls transcriptional dynamics during development. Science 375. doi:10.1126/SCIENCE.ABI7178.

Benko, S., Fantes, J. A., Amiel, J., Kleinjan, D. J., Thomas, S., Ramsay, J., et al. (2009). Highly conserved non-coding elements on either side of SOX9 associated with Pierre Robin sequence. Nat. Genet. 41, 359–364. doi:10.1038/ng.329.

Bergman, D. T., Jones, T. R., Liu, V., Ray, J., Jagoda, E., Siraj, L., et al. (2022). Compatibility rules of human enhancer and promoter sequences. Nature. doi:10.1038/S41586-022-04877-W.

Buecker, C., and Wysocka, J. (2012). Enhancers as information integration hubs in development: lessons from genomics. Trends Genet. 28, 276–284. doi:10.1016/j.tig.2012.02.008.

Bulger, M., and Groudine, M. (2010). Enhancers: The abundance and function of regulatory sequences beyond promoters. Dev. Biol. 339, 250–257. doi:10.1016/j.ydbio.2009.11.035.

Bulger, M., and Groudine, M. (2011). Functional and mechanistic diversity of distal transcription enhancers. Cell 144, 327–339. doi:10.1016/j.cell.2011.01.024.

Bult, C. J., Blake, J. A., Smith, C. L., Kadin, J. A., Richardson, J. E., Anagnostopoulos, A., et al. (2019). Mouse Genome Database (MGD) 2019. Nucleic Acids Res. 47, D801–D806. doi:10.1093/nar/gky1056.

Claringbould, A., and Zaugg, J. B. (2021). Enhancers in disease: molecular basis and emerging treatment strategies. Trends Mol. Med. 27, 1060–1073. doi:10.1016/J.MOLMED.2021.07.012/ATTACHMENT/84988D99-2316-4D3F-9DA1-26ADCA75651E/MMC1.XLSX.

Collins, R. L., Brand, H., Karczewski, K. J., Zhao, X., Alföldi, J., Francioli, L. C., et al. (2020). A structural variation reference for medical and population genetics. Nature 581, 444–451. doi:10.1038/s41586-020-2287-8.

Cox, J. J., Willatt, L., Homfray, T., and Woods, C. G. (2011). A SOX9 Duplication and Familial 46,XX Developmental Testicular Disorder. N. Engl. J. Med. 364, 91–93. doi:10.1056/nejmc1010311.

Creyghton, M. P., Cheng, A. W., Welstead, G. G., Kooistra, T., Carey, B. W., Steine, E. J., et al. (2010). Histone H3K27ac separates active from poised enhancers and predicts developmental state. Proc. Natl. Acad. Sci. U. S. A. 107, 21931–21936. doi:10.1073/PNAS.1016071107/-/DCSUPPLEMENTAL/PNAS.201016071SI.PDF.

D’haene, E., and Vergult, S. (2021). Interpreting the impact of noncoding structural variation in neurodevelopmental disorders. Genet. Med. 23, 34–46. doi:10.1038/s41436-020-00974-1.

Dixon, J. R., Selvaraj, S., Yue, F., Kim, A., Li, Y., Shen, Y., et al. (2012). Topological domains in mammalian genomes identified by analysis of chromatin interactions. Nature 485, 376–380. doi:10.1038/nature11082.

Dunham, I., Kundaje, A., Aldred, S. F., Collins, P. J., Davis, C. A., Doyle, F., et al. (2012). An integrated encyclopedia of DNA elements in the human genome. Nature 489, 57–74. doi:10.1038/nature11247.

Elgar, G., and Vavouri, T. (2008). Tuning in to the signals: noncoding sequence conservation in vertebrate genomes. Trends Genet. 24, 344–52. doi:10.1016/j.tig.2008.04.005.

Eraslan, G., Drokhlyansky, E., Anand, S., Fiskin, E., Subramanian, A., Slyper, M., et al. (2022). Single-nucleus cross-tissue molecular reference maps toward understanding disease gene function. Science (80-. ). 376. doi:10.1126/SCIENCE.ABL4290.

Federici, G., and Soddu, S. (2020). Variants of uncertain significance in the era of high-throughput genome sequencing: A lesson from breast and ovary cancers. J. Exp. Clin. Cancer Res. 39, 1–12. doi:10.1186/S13046-020-01554-6/FIGURES/1.

Feuk, L., Carson, A. R., and Scherer, S. W. (2006). Structural variation in the human genome. Nat. Rev. Genet. 7, 85–97. doi:10.1038/nrg1767.

Franke, M., Ibrahim, D. M., Andrey, G., Schwarzer, W., Heinrich, V., Schöpflin, R., et al. (2016). Formation of new chromatin domains determines pathogenicity of genomic duplications. Nature 538, 265–269. doi:10.1038/nature19800.

Fulco, C. P., Nasser, J., Jones, T. R., Munson, G., Bergman, D. T., Subramanian, V., et al. (2019). Activity-by-contact model of enhancer–promoter regulation from thousands of CRISPR perturbations. Nat. Genet. 51, 1664–1669. doi:10.1038/s41588-019-0538-0.

Galouzis, C. C., and Furlong, E. E. M. (2022). Regulating specificity in enhancer-promoter communication. Curr. Opin. Cell Biol. 75. doi:10.1016/J.CEB.2022.01.010.

Ganel, L., Abel, H. J., and Hall, I. M. (2017). SVScore: An impact prediction tool for structural variation. Bioinformatics 33, 1083–1085. doi:10.1093/bioinformatics/btw789.

Gerrard, D. T., Berry, A. A., Jennings, R. E., Birket, M. J., Zarrineh, P., Garstang, M. G., et al. (2020). Dynamic changes in the epigenomic landscape regulate human organogenesis and link to developmental disorders. Nat. Commun. 2020 111 11, 1–15. doi:10.1038/s41467-020-17305-2.

Gerrard, D. T., Berry, A. A., Jennings, R. E., Piper Hanley, K., Bobola, N., and Hanley, N. A. (2016). An integrative transcriptomic atlas of organogenesis in human embryos. Elife 5. doi:10.7554/eLife.15657.

Ghavi-Helm, Y. (2019). Functional consequences of chromosomal rearrangements on gene expression: not so deleterious after all? J. Mol. Biol. doi:10.1016/j.jmb.2019.09.010.

Hansen, B. H., Oerbeck, B., Skirbekk, B., Petrovski, B. É., and Kristensen, H. (2018). Neurodevelopmental disorders: prevalence and comorbidity in children referred to mental health services. Nord. J. Psychiatry 72, 285–291. doi:10.1080/08039488.2018.1444087.

Heintzman, N. D., Hon, G. C., Hawkins, R. D., Kheradpour, P., Stark, A., Harp, L. F., et al. (2009). Histone modifications at human enhancers reflect global cell-type-specific gene expression. Nature 459, 108–112. doi:10.1038/nature07829.

Hill, R. E., and Lettice, L. A. (2013). Alterations to the remote control of Shh gene expression cause congenital abnormalities. Philos. Trans. R. Soc. B Biol. Sci. 368, 20120357. doi:10.1098/RSTB.2012.0357.

Hertzberg, J., Mundlos, S., Vingron, M., and Gallone, G. (2022). TADA—a machine learning tool for functional annotation-based prioritisation of pathogenic CNVs. Genome Biol. 23, 1–21. doi:10.1186/S13059-022-02631-Z/FIGURES/4.

Ho, S. S., Urban, A. E., and Mills, R. E. (2020). Structural variation in the sequencing era. Nat. Rev. Genet. 21, 171–189. doi:10.1038/s41576-019-0180-9.

Home - MedGen - NCBI Available at: https://www.ncbi.nlm.nih.gov/medgen [Accessed May 20, 2022].

Ibn-Salem, J., Köhler, S., Love, M. I., Chung, H.-R., Huang, N., Hurles, M. E., et al. (2014). Deletions of chromosomal regulatory boundaries are associated with congenital disease. Genome Biol. 15, 423. doi:10.1186/s13059-014-0423-1.

Jeste, S. S. (2015). Neurodevelopmental behavioral and cognitive disorders. Contin. Lifelong Learn. Neurol. 21, 690–714. doi:10.1212/01.CON.0000466661.89908.3c.

Karczewski, K. J., Francioli, L. C., Tiao, G., Cummings, B. B., Alföldi, J., Wang, Q., et al. (2020). The mutational constraint spectrum quantified from variation in 141,456 humans. Nature 581, 434–443. doi:10.1038/s41586-020-2308-7.

Kirby, R. S. (2017). The prevalence of selected major birth defects in the United States. Semin. Perinatol. 41, 338–344. doi:10.1053/j.semperi.2017.07.004.

Klopocki, E., Ott, C. E., Benatar, N., Ullmann, R., Mundlos, S., and Lehmann, K. (2008). A microduplication of the long range SHH limb regulator (ZRS) is associated with triphalangeal thumb-polysyndactyly syndrome. J. Med. Genet. 45, 370–375. doi:10.1136/jmg.2007.055699.

Kraft, K., Magg, A., Heinrich, V., Riemenschneider, C., Schöpflin, R., Markowski, J., et al. (2019). Serial genomic inversions induce tissue-specific architectural stripes, gene misexpression and congenital malformations. Nat. Cell Biol. 21, 305–310. doi:10.1038/s41556-019-0273-x.

Krijger, P. H. L., and De Laat, W. (2016). Regulation of disease-associated gene expression in the 3D genome. Nat. Rev. Mol. Cell Biol. 17, 771–782. doi:10.1038/nrm.2016.138.

Krude, H., Mundlos, S., Øien, N. C., Opitz, R., and Schuelke, M. (2021). What can go wrong in the non-coding genome and how to interpret whole genome sequencing data. Medizinische Genet. 33, 121–131. doi:10.1515/medgen-2021-2071.

Kumakura, A., Takahashi, S., Okajima, K., and Hata, D. (2014). A haploinsufficiency of FOXG1 identified in a boy with congenital variant of Rett syndrome. Brain Dev. 36, 725–729. doi:10.1016/J.BRAINDEV.2013.09.006.

Landrum, M. J., Lee, J. M., Benson, M., Brown, G. R., Chao, C., Chitipiralla, S., et al. (2018). ClinVar: improving access to variant interpretations and supporting evidence. Nucleic Acids Res. 46, D1062–D1067. doi:10.1093/nar/gkx1153.

Lappalainen, I., Lopez, J., Skipper, L., Hefferon, T., Spalding, J. D., Garner, J., et al. (2013). DbVar and DGVa: Public archives for genomic structural variation. Nucleic Acids Res. 41. doi:10.1093/nar/gks1213.

Lappalainen, T., Scott, A. J., Brandt, M., and Hall, I. M. (2019). Genomic Analysis in the Age of Human Genome Sequencing. Cell 177, 70–84. doi:10.1016/j.cell.2019.02.032.

Laugsch, M., Bartusel, M., Rehimi, R., Alirzayeva, H., Karaolidou, A., Crispatzu, G., et al. (2019). Modeling the Pathological Long-Range Regulatory Effects of Human Structural Variation with Patient-Specific hiPSCs. Cell Stem Cell 24, 736–752.e12. doi:10.1016/j.stem.2019.03.004.

Lek, M., Karczewski, K. J., Minikel, E. V., Samocha, K. E., Banks, E., Fennell, T., et al. (2016). Analysis of protein-coding genetic variation in 60,706 humans. Nature 536, 285–291. doi:10.1038/nature19057.

Lessel, D., Gehbauer, C., Bramswig, N. C., Schluth-Bolard, C., Venkataramanappa, S., van Gassen, K. L. I., et al. (2018). BCL11B mutations in patients affected by a neurodevelopmental disorder with reduced type 2 innate lymphoid cells. Brain 141, 2299–2311. doi:10.1093/brain/awy173.

Lettice, L. A. (2003). A long-range Shh enhancer regulates expression in the developing limb and fin and is associated with preaxial polydactyly. Hum. Mol. Genet. 12, 1725–1735. doi:10.1093/hmg/ddg180.

Lettice, L. A., Daniels, S., Sweeney, E., Venkataraman, S., Devenney, P. S., Gautier, P., et al. (2011). Enhancer-adoption as a mechanism of human developmental disease. Hum. Mutat. 32, 1492–1499. doi:10.1002/humu.21615.

Lieberman-Aiden, E., van Berkum, N. L., Williams, L., Imakaev, M., Ragoczy, T., Telling, A., et al. (2009). Comprehensive mapping of long-range interactions reveals folding principles of the human genome. Science 326, 289–93. doi:10.1126/science.1181369.

Lipton, Z. C. (2016). The Mythos of Model Interpretability. Commun. ACM 61, 35–43. doi:10.48550/arxiv.1606.03490.

Lohan, S., Spielmann, M., Doelken, S. C., Flöttmann, R., Muhammad, F., Baig, S. M., et al. (2014). Microduplications encompassing the sonic hedgehog limb enhancer ZRS are associated with haas-type polysyndactyly and Laurin-Sandrow syndrome. Clin. Genet. 86, 318–325. doi:10.1111/cge.12352.

Long, H. K., Osterwalder, M., Welsh, I. C., Hansen, K., Davies, J. O. J., Liu, Y. E., et al. (2020). Loss of Extreme Long-Range Enhancers in Human Neural Crest Drives a Craniofacial Disorder. Cell Stem Cell 27, 765–783.e14. doi:10.1016/j.stem.2020.09.001.

Louden, D. N. (2020). MedGen: NCBI’s Portal to Information on Medical Conditions with a Genetic Component. https://doi.org/10.1080/02763869.2020.1726152 39, 183–191. doi:10.1080/02763869.2020.1726152.

Lupiáñez, D. G., Kraft, K., Heinrich, V., Krawitz, P., Brancati, F., Klopocki, E., et al. (2015). Disruptions of Topological Chromatin Domains Cause Pathogenic Rewiring of Gene-Enhancer Interactions. Cell 161, 1012–1025. doi:10.1016/j.cell.2015.04.004.

Lupiáñez, D. G., Spielmann, M., and Mundlos, S. (2016). Breaking TADs: How Alterations of Chromatin Domains Result in Disease. Trends Genet. 32, 225–237. doi:10.1016/j.tig.2016.01.003.

McLaren, W., Gil, L., Hunt, S. E., Riat, H. S., Ritchie, G. R. S., Thormann, A., et al. (2016). The Ensembl Variant Effect Predictor. Genome Biol. 17, 122. doi:10.1186/s13059-016-0974-4.

Mehrjouy, M. M., Fonseca, A. C. S., Ehmke, N., Paskulin, G., Novelli, A., Benedicenti, F., et al. (2018). Regulatory variants of FOXG1 in the context of its topological domain organisation/631/208/200 /631/208/1516 article. Eur. J. Hum. Genet. 26, 186–196. doi:10.1038/s41431-017-0011-4.

Middelkamp, S., Vlaar, J. M., Giltay, J., Korzelius, J., Besselink, N., Boymans, S., et al. (2019). Prioritization of genes driving congenital phenotypes of patients with de novo genomic structural variants. Genome Med. 11, 79. doi:10.1186/s13073-019-0692-0.

Milunsky, J. M., Maher, T. A., Zhao, G., Roberts, A. E., Stalker, H. J., Zori, R. T., et al. (2008). TFAP2A Mutations Result in Branchio-Oculo-Facial Syndrome. Am. J. Hum. Genet. 82, 1171–1177. doi:10.1016/j.ajhg.2008.03.005.

Nanni, L., Ming, J. E., Bocian, M., Steinhaus, K., Bianchi, D. W., De Die-Smulders, C., et al. (1999). The mutational spectrum of the sonic hedgehog gene in holoprosencephaly: SHH mutations cause a significant proportion of autosomal dominant holoprosencephaly. Hum. Mol. Genet. 8, 2479–2488. doi:10.1093/HMG/8.13.2479.

Nichols, J. A., Herbert Chan, H. W., and Baker, M. A. B. (2019). Machine learning: applications of artificial intelligence to imaging and diagnosis. Biophys. Rev. 11, 111. doi:10.1007/S12551-018-0449-9.

Nora, E. P., Dekker, J., and Heard, E. (2013). Segmental folding of chromosomes: A basis for structural and regulatory chromosomal neighborhoods? BioEssays 35, 818–828. doi:10.1002/bies.201300040.

Ong, C.-T., and Corces, V. G. (2011). Enhancer function: new insights into the regulation of tissue-specific gene expression. Nat. Rev. Genet. 12, 283–293. doi:10.1038/nrg2957.

Pachano, T., Sánchez-Gaya, V., Ealo, T., Mariner-Faulí, M., Bleckwehl, T., Asenjo, H. G., et al. (2021). Orphan CpG islands amplify poised enhancer regulatory activity and determine target gene responsiveness. Nat. Genet. 53, 1036–1049. doi:10.1038/s41588-021-00888-x.

Pereira, R., Halford, K., Sokolov, B. P., Khillan, J. S., and Prockop, D. J. (1994). Phenotypic variability and incomplete penetrance of spontaneous fractures in an inbred strain of transgenic mice expressing a mutated collagen gene (COL1A1). J. Clin. Invest. 93, 1765–1769. doi:10.1172/JCI117161.

Poszewiecka, B., Pienkowski, V. M., Nowosad, K., Erôme, J., Robin, D., Gogolewski, K., et al. (2022). TADeus2: a web server facilitating the clinical diagnosis by pathogenicity assessment of structural variations disarranging 3D chromatin structure. Nucleic Acids Res. 1, 13–14. doi:10.1093/NAR/GKAC318.

Quinlan, A. R., and Hall, I. M. (2010). BEDTools: a flexible suite of utilities for comparing genomic features. Bioinformatics 26, 841–842. doi:10.1093/bioinformatics/btq033.

Quinn, T. P., Jacobs, S., Senadeera, M., Le, V., and Coghlan, S. (2022). The three ghosts of medical AI: Can the black-box present deliver? Artif. Intell. Med. 124, 102158. doi:10.1016/J.ARTMED.2021.102158.

Rada-Iglesias, A., Bajpai, R., Swigut, T., Brugmann, S. A., Flynn, R. A., and Wysocka, J. (2011). A unique chromatin signature uncovers early developmental enhancers in humans. Nature 470, 279–283. doi:10.1038/nature09692.

Rada-Iglesias, A., and Wysocka, J. (2011). Epigenomics of human embryonic stem cells and induced pluripotent stem cells: insights into pluripotency and implications for disease. Genome Med. 3, 36. doi:10.1186/gm252.

Rainger, J. K., Bhatia, S., Bengani, H., Gautier, P., Rainger, J., Pearson, M., et al. (2014). Disruption of SATB2 or its long-range cis-regulation by SOX9 causes a syndromic form of Pierre Robin sequence. Hum. Mol. Genet. 23, 2569–2579. doi:10.1093/hmg/ddt647.

Rao, S. S. P., Huntley, M. H., Durand, N. C., Stamenova, E. K., Bochkov, I. D., Robinson, J. T., et al. (2014). A 3D Map of the Human Genome at Kilobase Resolution Reveals Principles of Chromatin Looping. Cell 159, 1665–1680. doi:10.1016/j.cell.2014.11.021.

Real, F. M., Haas, S. A., Franchini, P., Xiong, P., Simakov, O., Kuhl, H., et al. (2020). The mole genome reveals regulatory rearrangements associated with adaptive intersexuality. Science 370, 208–214. doi:10.1126/SCIENCE.AAZ2582.

Reddy, S. (2022). Explainability and artificial intelligence in medicine. doi:10.1016/S2589-7500(22)00029-2.

Redin, C., Brand, H., Collins, R. L., Kammin, T., Mitchell, E., Hodge, J. C., et al. (2017). The genomic landscape of balanced cytogenetic abnormalities associated with human congenital anomalies. Nat. Genet. 49, 36–45. doi:10.1038/ng.3720.

Rehimi, R., Nikolic, M., Cruz-Molina, S., Tebartz, C., Frommolt, P., Mahabir, E., et al. (2016). Epigenomics-Based Identification of Major Cell Identity Regulators within Heterogeneous Cell Populations. Cell Rep. 17, 3062–3076. doi:10.1016/J.CELREP.2016.11.046.

Richards, S., Aziz, N., Bale, S., Bick, D., Das, S., Gastier-Foster, J., et al. (2015). Standards and guidelines for the interpretation of sequence variants: a joint consensus recommendation of the American College of Medical Genetics and Genomics and the Association for Molecular Pathology. Genet. Med. 17, 405–424. doi:10.1038/GIM.2015.30.

Ringel, A. R., Szabo, Q., Chiariello, A. M., Chudzik, K., Schöpflin, R., Rothe, P., et al. (2021). Promoter repression and 3D-restructuring resolves divergent developmental gene expression in TADs. bioRxiv, 2021.10.08.463672. doi:10.1101/2021.10.08.463672.

Rodriguez-Revenga, L., Mila, M., Rosenberg, C., Lamb, A., and Lee, C. (2007). Structural variation in the human genome: The impact of copy number variants on clinical diagnosis. Genet. Med. 9, 600–606. doi:10.1097/GIM.0b013e318149e1e3.

Sagai, T., Hosoya, M., Mizushina, Y., Tamura, M., and Shiroishi, T. (2005). Elimination of a long-range cis-regulatory module causes complete loss of limb-specific Shh expression and truncation of the mouse limb. Development 132, 797–803. doi:10.1242/dev.01613.

Samanta, D. (2020). PCDH19-Related Epilepsy Syndrome: A Comprehensive Clinical Review. Pediatr. Neurol. 105, 3–9. doi:10.1016/j.pediatrneurol.2019.10.009.

Sánchez-Gaya, V., Mariner-Faulí, M., and Rada-Iglesias, A. (2020). Rare or Overlooked? Structural Disruption of Regulatory Domains in Human Neurocristopathies. Front. Genet. 11, 688. doi:10.3389/fgene.2020.00688.

Sanyal, A., Lajoie, B. R., Jain, G., and Dekker, J. (2012). The long-range interaction landscape of gene promoters. Nature 489, 109–113. doi:10.1038/nature11279.

Satterlee, J. S., Chadwick, L. H., Tyson, F. L., McAllister, K., Beaver, J., Birnbaum, L., et al. (2019). The NIH Common Fund/Roadmap Epigenomics Program: Successes of a comprehensive consortium. Sci. Adv. 5. doi:10.1126/SCIADV.AAW6507/ASSET/53CF23FC-E72F-490B-BD65-90BE29A0B9AE/ASSETS/GRAPHIC/AAW6507-F2.JPEG.

Sharo, A. G., Hu, Z., Sunyaev, S. R., and Brenner, S. E. (2022). StrVCTVRE: A supervised learning method to predict the pathogenicity of human genome structural variants. Am. J. Hum. Genet. 109, 195–209. doi:10.1016/j.ajhg.2021.12.007.

Shim, W. J., Sinniah, E., Xu, J., Vitrinel, B., Alexanian, M., Andreoletti, G., et al. (2020). Conserved Epigenetic Regulatory Logic Infers Genes Governing Cell Identity. Cell Syst. 11, 625–639.e13. doi:10.1016/j.cels.2020.11.001.

Smith, L., Singhal, N., El Achkar, C. M., Truglio, G., Rosen Sheidley, B., Sullivan, J., et al. (2018). PCDH19-related epilepsy is associated with a broad neurodevelopmental spectrum. Epilepsia 59, 679–689. doi:10.1111/epi.14003.

Spielmann, M., and Mundlos, S. (2016). Looking beyond the genes: the role of non-coding variants in human disease. Hum. Mol. Genet. 25, R157–R165. doi:10.1093/HMG/DDW205.

Spielmann, M., Lupiáñez, D. G., and Mundlos, S. (2018). Structural variation in the 3D genome. Nat. Rev. Genet. 19, 453–467. doi:10.1038/s41576-018-0007-0.

Spector, J. D., and Wiita, A. P. (2019). ClinTAD: a tool for copy number variant interpretation in the context of topologically associated domains. J. Hum. Genet. 64, 437–443. doi:10.1038/s10038-019-0573-9.

Stankiewicz, P., and Lupski, J. R. (2010). Structural variation in the human genome and its role in disease. Annu. Rev. Med. 61, 437–455. doi:10.1146/annurev-med-100708-204735.

Symonds, J. D., Zuberi, S. M., Stewart, K., McLellan, A., O’Regan, M., MacLeod, S., et al. (2019). Incidence and phenotypes of childhood-onset genetic epilepsies: A prospective population-based national cohort. Brain 142, 2303–2318. doi:10.1093/brain/awz195.

Tabula Sapiens Consortium*, Jones, R. C., Karkanias, J., Krasnow, M. A., Pisco, A. O., Quake, S. R., et al. (2022). The Tabula Sapiens: A multiple-organ, single-cell transcriptomic atlas of humans. Science 376, eabl4896. doi:10.1126/SCIENCE.ABL4896/SUPPL_FILE/SCIENCE.ABL4896_MDAR_REPRODUCIBILITY_CHECKLIST.PDF.

Thorwarth, A., Sarah, S. H., Schrumpf, P., Müller, I., Jyrch, S., Dame, C., et al. (2014). Comprehensive genotyping and clinical characterisation reveal 27 novel NKX2-1 mutations and expand the phenotypic spectrum. J. Med. Genet. 51, 375–387. doi:10.1136/jmedgenet-2013-102248.

Tocco, C., Bertacchi, M., and Studer, M. (2021). Structural and Functional Aspects of the Neurodevelopmental Gene NR2F1: From Animal Models to Human Pathology. Front. Mol. Neurosci. 14, 279. doi:10.3389/FNMOL.2021.767965/BIBTEX.

Trainor, P. A. (2010). Craniofacial birth defects: The role of neural crest cells in the etiology and pathogenesis of Treacher Collins syndrome and the potential for prevention. Am. J. Med. Genet. Part A 152A, 2984–2994. doi:10.1002/ajmg.a.33454.

Vandermeer, J. E., Smith, R. P., Jones, S. L., and Ahituv, N. (2014). Genome-wide identification of signaling center enhancers in the developing limb. Dev. 141, 4194–4198. doi:10.1242/dev.110965.

Vellido, A. (2020). The importance of interpretability and visualization in machine learning for applications in medicine and health care. Neural Comput. Appl. 32, 18069–18083. doi:10.1007/S00521-019-04051-W/TABLES/2.

Wittkopp, P. J., and Kalay, G. (2012). Cis-regulatory elements: molecular mechanisms and evolutionary processes underlying divergence. Nat. Rev. Genet. 13, 59–69. doi:10.1038/nrg3095.

Wray, G. A. (2007). The evolutionary significance of cis-regulatory mutations. Nat. Rev. Genet. 8, 206–216. doi:10.1038/nrg2063.

Wu, W., He, J., and Shao, X. (2020). Incidence and mortality trend of congenital heart disease at the global, regional, and national level, 1990-2017. Med. (United States) 99. doi:10.1097/MD.0000000000020593.

Zhang, F., and Lupski, J. R. (2015). Non-coding genetic variants in human disease. doi:10.1093/hmg/ddv259.

Zhang, Z., and Zhao, Y. (2022). Progress on the roles of MEF2C in neuropsychiatric diseases. Mol. Brain 15. doi:10.1186/S13041-021-00892-6.

Zhong, S., Zhang, S., Fan, X., Wu, Q., Yan, L., Dong, J., et al. (2018). A single-cell RNA-seq survey of the developmental landscape of the human prefrontal cortex. Nature 555, 524–528. doi:10.1038/nature25980.

Zhu, Y., Tazearslan, C., and Suh, Y. (2017). Challenges and progress in interpretation of non-coding genetic variants associated with human disease. Exp. Biol. Med. 242, 1325–1334. doi:10.1177/1535370217713750.

Zuin, J., Roth, G., Zhan, Y., Cramard, J., Redolfi, J., Piskadlo, E., et al. (2022). Nonlinear control of transcription through enhancer-promoter interactions. Nature 604, 571–577. doi:10.1038/S41586-022-04570-Y.

