## Supplementary Data 1 for "POSTRE: a tool to predict the pathological effects of human structural variants": Supplementary_Data_1.html

xml version="1.0" encoding="UTF-8"?
Cell Types/Tissues per Phenotype 

|  |  |  |  |  |  |  |  |  |
| --- | --- | --- | --- | --- | --- | --- | --- | --- |
| Phenotype | Cell Types/Tissues | Cell Types/Tissues additional information | Chip-Seq Source for enhancer annotation | Enhancer annotation (based on Chip-Seq), Main Procedure | RNA-Seq Source | Gene Expression quantification main procedure | HiC – TAD maps Source | TAD maps obtaining main procedure |
| Cardiovascular | Day 5 | Day 5 cardiac mesodermal cells, from 80 days ESC differentiation towards ventricular cardiomocytes  For + info: https://www.nature.com/articles/s41588-019-0479-7 | https://doi.org/10.1038/s41588-019-0479-7 | H3K27ac peak with distance > 10Kb to protein coding TSS  (Peaks were called after reads mapping with Bowtie2 with Macs2) | https://doi.org/10.1038/s41588-019-0479-7 | FPKMs directly obtained from source | https://doi.org/10.1038/s41588-019-0479-7 | TAD map generated upon processing Heart day 5 HiC data.  TADs were called with the usage of the tool DomainCaller by considering the 50kb contact matrices in the .hic files |
|  | Day 7 | Day 7 cardiac progenitors, from 80 days ESC differentiation towards ventricular cardiomocytes  For + info: https://www.nature.com/articles/s41588-019-0479-7 | https://doi.org/10.1038/s41588-019-0479-7 | H3K27ac peak with distance > 10Kb to protein coding TSS  (Peaks were called after reads mapping with Bowtie2 with Macs2) | https://doi.org/10.1038/s41588-019-0479-7 | FPKMs directly obtained from source | https://doi.org/10.1038/s41588-019-0479-7 | TAD map generated upon processing Heart day 7 HiC data.  TADs were called with the usage of the tool DomainCaller by considering the 50kb contact matrices in the .hic files |
|  | Day 15 | Day 15 primitive cardiomyocytes, from 80 days ESC differentiation towards ventricular cardiomocytes  For + info: https://www.nature.com/articles/s41588-019-0479-7 | https://doi.org/10.1038/s41588-019-0479-7 | H3K27ac peak with distance > 10Kb to protein coding TSS  (Peaks were called after reads mapping with Bowtie2 with Macs2) | https://doi.org/10.1038/s41588-019-0479-7 | FPKMs directly obtained from source | https://doi.org/10.1038/s41588-019-0479-7 | TAD map generated upon processing Heart day 15 HiC data.  TADs were called with the usage of the tool DomainCaller by considering the 50kb contact matrices in the .hic files |
|  | Day 80 | Day 80 ventricular cardiomiocytes, from 80 days ESC differentiation towards ventricular cardiomocytes  For + info: https://www.nature.com/articles/s41588-019-0479-7 | https://doi.org/10.1038/s41588-019-0479-7 | H3K27ac peak with distance > 10Kb to protein coding TSS  (Peaks were called after reads mapping with Bowtie2 with Macs2) | https://doi.org/10.1038/s41588-019-0479-7 | FPKMs directly obtained from source | https://doi.org/10.1038/s41588-019-0479-7 | TAD map generated upon processing Heart day 80 HiC data.  TADs were called with the usage of the tool DomainCaller by considering the 50kb contact matrices in the .hic files |
| Head-Neck | Neural Crest Early | Neural Crest Data of hNCC differentiation  This NC data corresponds with an early and heterogeneous stage of NC differentiation  For +info: https://pubmed.ncbi.nlm.nih.gov/29719267/ | https://www.ncbi.nlm.nih.gov/geo/query/acc.cgi?acc=GSE28876 | p300 peak, intersecting H3K27ac peak, with distance > 10Kb to protein coding TSS  (Peaks were called upon reads mapping with Bowtie2 with Macs2) | https://www.ncbi.nlm.nih.gov/geo/query/acc.cgi?acc=GSE28876 | FPKMs obtained upon reads mapping with Tophat2 and gene expression quantification with Cufflinks | 3D genome browser: http://3dgenome.fsm.northwestern.edu/  TAD maps: http://3dgenome.fsm.northwestern.edu/downloads/hg19.TADs.zip | ESC TAD map directly obtained from: http://3dgenome.fsm.northwestern.edu/downloads/hg19.TADs.zip |
|  | Neural Crest Late | Neural Crest Data of hNCC differentiation  P4 differentiation stage, it corresponds with a later and more homogeneous stage of NC differentation in comparison with Neural Crest Early  For +info: https://www.ncbi.nlm.nih.gov/geo/query/acc.cgi?acc=GSE70751 | https://www.ncbi.nlm.nih.gov/geo/query/acc.cgi?acc=GSE70751 | p300 peak, intersecting H3K27ac peak, with distance > 10Kb to protein coding TSS  (Peaks were called upon reads mapping with Bowtie2 with Macs2) | https://www.ncbi.nlm.nih.gov/geo/query/acc.cgi?acc=GSE70751 | FPKMs obtained upon reads mapping with Tophat2 and  gene expression quantification with Cufflinks | 3D genome browser: http://3dgenome.fsm.northwestern.edu/  TAD maps: http://3dgenome.fsm.northwestern.edu/downloads/hg19.TADs.zip | ESC TAD map directly obtained from: http://3dgenome.fsm.northwestern.edu/downloads/hg19.TADs.zip |
|  | PalateCS20 | Embryonic Palate Carnagie Stage 20  For +info: https://pubmed.ncbi.nlm.nih.gov/29719267/ | https://pubmed.ncbi.nlm.nih.gov/29719267/ | H3K27ac peak with distance > 10Kb to protein coding TSS  (Peaks were called with Macs2 using the provided alignment files (tagAlign)) | https://elifesciences.org/articles/15657 | To obtain the FPKMs, first, the bedfiles containing the reads mapping coordinates (.unique.bed12) for Palate1 sample were converted into bam files with bedToBam tool.  Next FPKMs were computed with Cufflinks. | 3D genome browser: http://3dgenome.fsm.northwestern.edu/  TAD maps: http://3dgenome.fsm.northwestern.edu/downloads/hg19.TADs.zip | ESC TAD map directly obtained from: http://3dgenome.fsm.northwestern.edu/downloads/hg19.TADs.zip |
| Limbs | EmbryonicLimb1 | Corresponds with embryonic Lower Limb data  For +info: https://www.nature.com/articles/s41467-020-17305-2 | https://www.nature.com/articles/s41467-020-17305-2 | H3K27ac peak with distance > 10Kb to protein coding TSS  (Peaks were called after reads mapping with Bowtie2 with Macs2) | https://elifesciences.org/articles/15657 | To obtain the FPKMs, first, the bedfiles containing the reads mapping coordinates (.unique.bed12) for Lower Limb samples were converted into bam files with bedToBam tool.  Next FPKMs were computed with Cufflinks. | 3D genome browser: http://3dgenome.fsm.northwestern.edu/  TAD maps: http://3dgenome.fsm.northwestern.edu/downloads/hg19.TADs.zip | ESC TAD map directly obtained from: http://3dgenome.fsm.northwestern.edu/downloads/hg19.TADs.zip |
|  | EmbryonicLimb2 | Corresponds with embryonic Upper Limb data  For +info: https://www.nature.com/articles/s41467-020-17305-2 | https://www.nature.com/articles/s41467-020-17305-2 | H3K27ac peak with distance > 10Kb to protein coding TSS  (Peaks were called after reads mapping with Bowtie2 with Macs2) | https://elifesciences.org/articles/15657 | To obtain the FPKMs, first, the bedfiles containing the reads mapping coordinates (.unique.bed12) for Upper Limb samples were converted into bam files with bedToBam tool.  Next FPKMs were computed with Cufflinks. | 3D genome browser: http://3dgenome.fsm.northwestern.edu/  TAD maps: http://3dgenome.fsm.northwestern.edu/downloads/hg19.TADs.zip | ESC TAD map directly obtained from: http://3dgenome.fsm.northwestern.edu/downloads/hg19.TADs.zip |
| Neurodevelopmental | PfcGw15 | Brain prefrontal cortex Gestation Week 15  For +info: https://www.ncbi.nlm.nih.gov/geo/query/acc.cgi?acc=GSE149268 | https://www.ncbi.nlm.nih.gov/geo/query/acc.cgi?acc=GSE149268 | H3K27ac peak with distance > 10Kb to protein coding TSS  (Peaks were directly obtained from source) | From database: https://www.brainspan.org/static/home | Expression values obtained from file: summarized to genes file  Given that gestational age is 2 weeks longer than conceptional age, post conception week 13 (pcw13) samples information was considered to match with chip-seq gestation week 15 data  Expression value computed as the average of the pcw13 samples: VFC (ventrolateral prefrontal cortex), MFC[anterior (rostral) cingulate (medial prefrontal) cortex] and DFC(dorsolateral prefrontal cortex) | https://doi.org/10.1016/j.celrep.2016.10.061 | Brain Prefrontal Cortex TAD map, generated from CO (prefrontal cortex) boundary map provided in the referenced source.  The Excelfile containing the boundary maps can be found here:  https://www.cell.com/cms/10.1016/j.celrep.2016.10.061/attachment/c3954472-3378-4fd7-aa33-9a2d085f7bbd/mmc4.xlsx |
|  | PfcGw18 | Brain prefrontal cortex Gestation Week 18  For +info: https://www.ncbi.nlm.nih.gov/geo/query/acc.cgi?acc=GSE149268 | https://www.ncbi.nlm.nih.gov/geo/query/acc.cgi?acc=GSE149268 | H3K27ac peak with distance > 10Kb to protein coding TSS  (Peaks were directly obtained from source) | From database: https://www.brainspan.org/static/home | Expression values obtained from file: summarized to genes file  Given that gestational age is 2 weeks longer than conceptional age, post conception week 16 (pcw16) samples information was considered to match with chip-seq gestation week 18 data  Expression value computed as the average of the pcw16 samples: VFC (ventrolateral prefrontal cortex), MFC[anterior (rostral) cingulate (medial prefrontal) cortex] and DFC(dorsolateral prefrontal cortex) | https://doi.org/10.1016/j.celrep.2016.10.061 | Brain Prefrontal Cortex TAD map, generated from CO (prefrontal cortex) boundary map provided in the referenced source.  The Excelfile containing the boundary maps can be found here:  https://www.cell.com/cms/10.1016/j.celrep.2016.10.061/attachment/c3954472-3378-4fd7-aa33-9a2d085f7bbd/mmc4.xlsx |
